# Unveiling correlational nexus among environment, gut microbiota, and personality traits in the Iberian Lynx (*Lynx pardinus*)

**DOI:** 10.64898/2026.06.25.734547

**Authors:** Agustín Carbajo-Usano, Xavier Triadó-Margarit, Antonio Rivas, Arnau Vendrell, Irene Gutierrez, Emilio O Casamayor

## Abstract

The gut microbiome is increasingly recognized as a pivotal modulator of animal behaviour, yet its influence on wild fauna remains largely unexplored. We investigated the correlational relationship between gut microbiota, behavioural phenotypes, and management practices in 26 captive endangered Iberian lynxs (*Lynx pardinus*) maintained within the *ex-situ* Iberian breeding program facilities, in two geographically distant stations in SW Spain. Behavioural observations were intensively recorded over two years, and three personality profiles emerged, i.e., (i) anomalous (with the highest frequencies for stereotypies), and (ii) sedentary and (iii) active (with the highest frequencies for sedentarism and for locomotion and surveillance, respectively). Fecal samples were analyzed for biweekly periods by 16S rRNA gene amplicon sequencing to profile bacterial composition and predicted functional pathways, and significant associations were found for each of the behavioural phenotypes. Both breeding station and local environment influenced gut microbial communities and personality profiles, underscoring the influence of management practices and local habitat in shaping the microbiome–behaviour nexus. Specific bacterial taxa and metabolic pathways were consistently associated with each behavioural phenotype, suggesting that microbial fecal signatures could serve as non-invasive biomarkers for individual personality monitoring. This work constitutes the first comprehensive, multi-layered examination of the interplay among behaviour, gut microbiota, and environmental factors in a large, wild carnivore. This integrative approach may help conservation programmes to optimize management decisions and improve reintroduction success.

## INTRODUCTION

The Iberian lynx (Lynx pardinus) is one of the most endangered felids in the world. Its decline has been driven by habitat loss and fragmentation, prey scarcity, vehicle collisions, and illegal hunting (Vargas et al., 2008). Since the implementation of the Iberian Lynx Ex situ Conservation Programme, captive breeding has played a central role in maintaining a genetically viable population while supporting reintroduction efforts. This programme provides a unique opportunity to investigate ecological and behavioural processes under controlled conditions which are relevant for conservation practices. In recent years, the Iberian lynx has been the focus of a wide array of studies, given its unique status and symbolic importance. Some of them have primarily examined movement patterns and exploratory behaviour (Rueda et al., 2021; Cisneros-Araujo et al., 2025; Goicolea et al., 2021), and a qualitative personality study was previously carried out, highlighting the importance of taking into account personality traits to improve breeding management and reintroduction success in the Iberian lynx (Úbeda et al., 2021).

Animal personality is usually understood as individual differences in behavioural tendencies that are present across time and contexts (Réale et al. 2007). The importance of studying individual personality and behaviour for conservation and reintroduction actions has been recently highlighted (Joseph et al., 2019), and ethological studies are considered before reintroducing species due to the implications for colonization and dispersal success. However, ethological studies are very difficult to carry out with wild fauna and most of the studies have been focused on small animals (Erica et al., 2022; Cheng, Jiming & He, 2023). Stereotypic behaviours, commonly observed in captive animals, are widely recognised as indicators of stress or suboptimal environmental conditions. Their occurrence is strongly associated with captive rearing (Hediger et al., 1950; Vestergaard et al., 1981), and represents a global challenge for species reintroduction programmes, potentially compromising adaptive success in the wild (Sohel Khan et al., 2022). Animal behaviour is not solely determined by intrinsic individual traits, but is it also shaped by the surrounding environmental factors (e.g., habitat structural complexity, vegetation cover…) as they can influence the expression of natural behaviours, as well as animal welfare and stress levels (Alejandro, J., 2022). Climatic conditions and management practices in captivity may further modulate these effects (Deborah L., 2009).

In addition to the factors mentioned above, increasing lines of evidence indicate that internal biological factors also contribute to modulate behavioural variation. Among these, the gut microbiota has emerged as a key regulator of host physiology, metabolism, immune function, and behaviour (Kuziel et al., 2022; Sekirov et al., 2010; Sommer et al., 2013). Gut microbial communities are shaped by multiple host and environment related factors, including diet, physiological status, stress, disease, geographical location, and life stage (Chang et al., 2019). In turn, the gut microbiota communicates bidirectionally with the central nervous system through neuroendocrine, neuroimmune, and neural pathways, collectively known as the gut-brain axis, influencing stress responsiveness, cognition, emotional regulation, and behavioural expression (Petra et al., 2015; Naufel et al., 2023). Environmental conditions may further influence these host–microbiota interactions by modifying resource availability, microbial exposure, physiological stress, and activity patterns. Consequently, behaviour, gut microbiota, and environmental conditions should be regarded as components of an integrated ecological system, where changes in one component may propagate through the others. Understanding behavioural variation therefore requires considering these interacting processes as a whole.

The Iberian lynx ex situ conservation programme provides an exceptional opportunity to address these interactions. Individuals are monitored longitudinally under standardized management while being housed in geographically separated breeding centres that differ in environmental characteristics, allowing detailed behavioural observations, frequent faecal sampling, and environmental assessment under realistic conservation conditions. Taking advantage of this unique framework, we aimed to simultaneously characterize personality profiles and gut microbiota composition in captive Iberian lynxes, evaluate how environmental conditions influence both components, and identify microbial signatures associated with behavioural phenotypes and management practices.

## MATERIAL Y METODOS

### 2.1. Source and selection of Iberian Lynx individuals

The samples were obtained from individuals of the Iberian Lynx Conservation Breeding Program (https://www.lynxexsitu.es/), a successful ex-situ conservation initiative designed to both (i) maintain a genetically and demographically managed captive population of *Lynx pardinus* distributed in several breeding centres of the Iberian Peninsula (https://www.lynxexsitu.es/programa-en.php?sec=centro) and (ii) create new reintroduced free-ranging populations in Spain and Portugal (Vargas et al. 2008 DOI:10.1111/j.1748-1090.2007.00036.x). The different breeding centers follow a common standardized official management protocol (https://www.lynxexsitu.es/seccion.php?secc=DOCUMENTOS&tipo=10), and the breeding individuals had a weekly fixed diet of live rabbit (x3), dead rabbit (x2), veal or poultry (x1) and fasting (x1). Captive individuals are usually kept alone in large fenced areas and are constantly recorded by video cameras 24h per day all along the year. Each individual is surveyed for 10 seconds every hour by trained volunteer staff, and the gathered behaviour is manually noted and classified in a common standardized ethogram (see below). We studied 26 individuals (Table 1) belonging to two breeding centres located in south-western Spain, i.e., 9 individuals (4 males and 5 females) from “Acebuche” station in the locality of Matalascañas near the Doñana National Park (Huelva, Andalusia), and 17 individuals (9 males and 8 females) from “Granadilla”, near the locality of Zarza de Granadilla (Cáceres, Extremadura). The individuals were studied between January 2018 and September 2020 in Acebuche, and between November 2021 to November 2023 in Granadilla. Publicly available environmental data for the selected sites (i.e., 10 m wind u-and v-components, 2 m dewpoint temperature, 2 m temperature, evaporation, total cloud cover, vegetation cover and total precipitation) were obtained along the study periods from the ERA5 dataset of the European Centre for Medium-Range Weather Forecasts (Copernicus Climate Change Service). Differences between the two breeding stations for each environmental variable was statistically tested by hypothesis contrast tests (t.test function of stats package version 3.6.2).

**TABLE 1.**
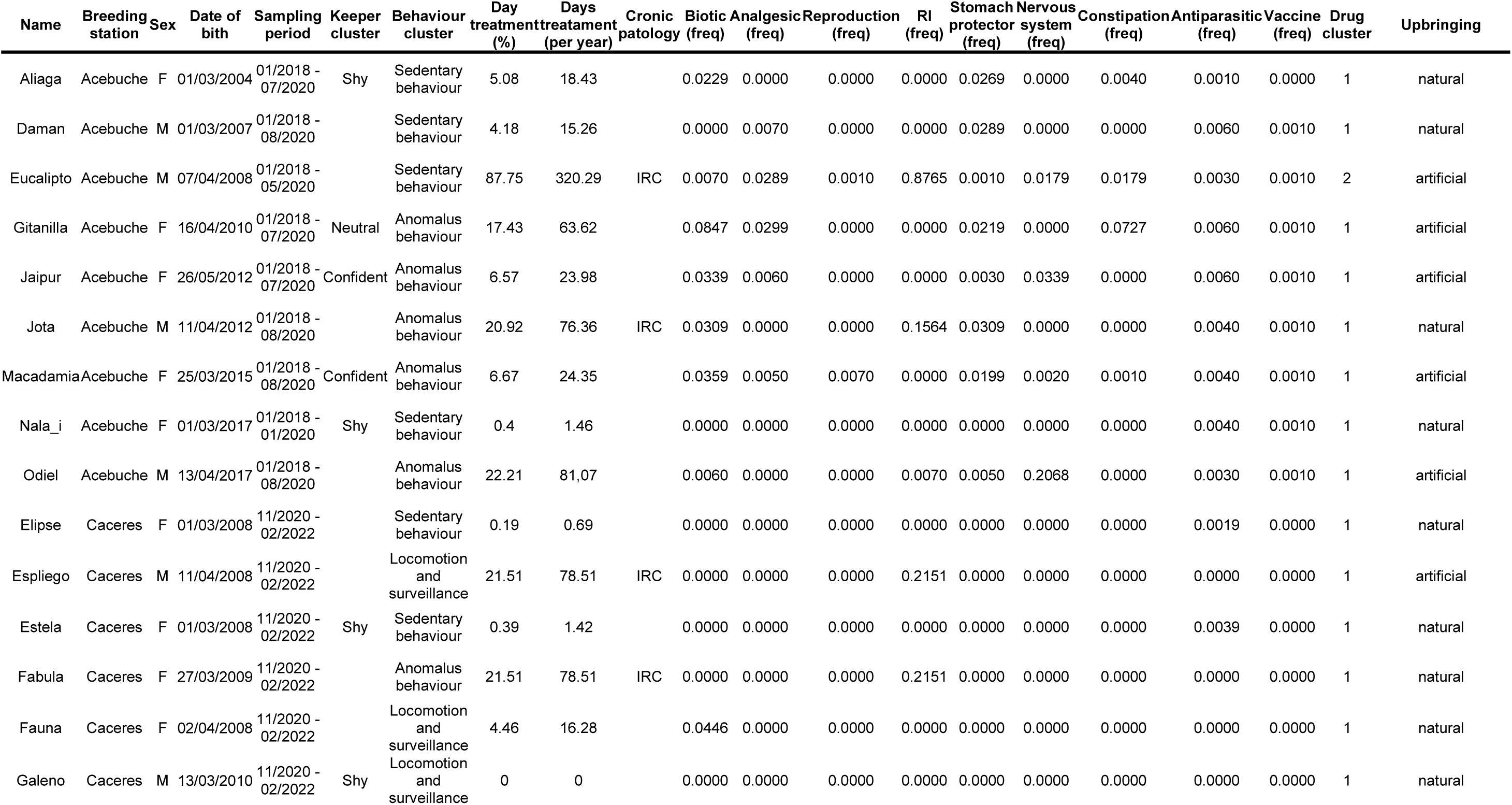
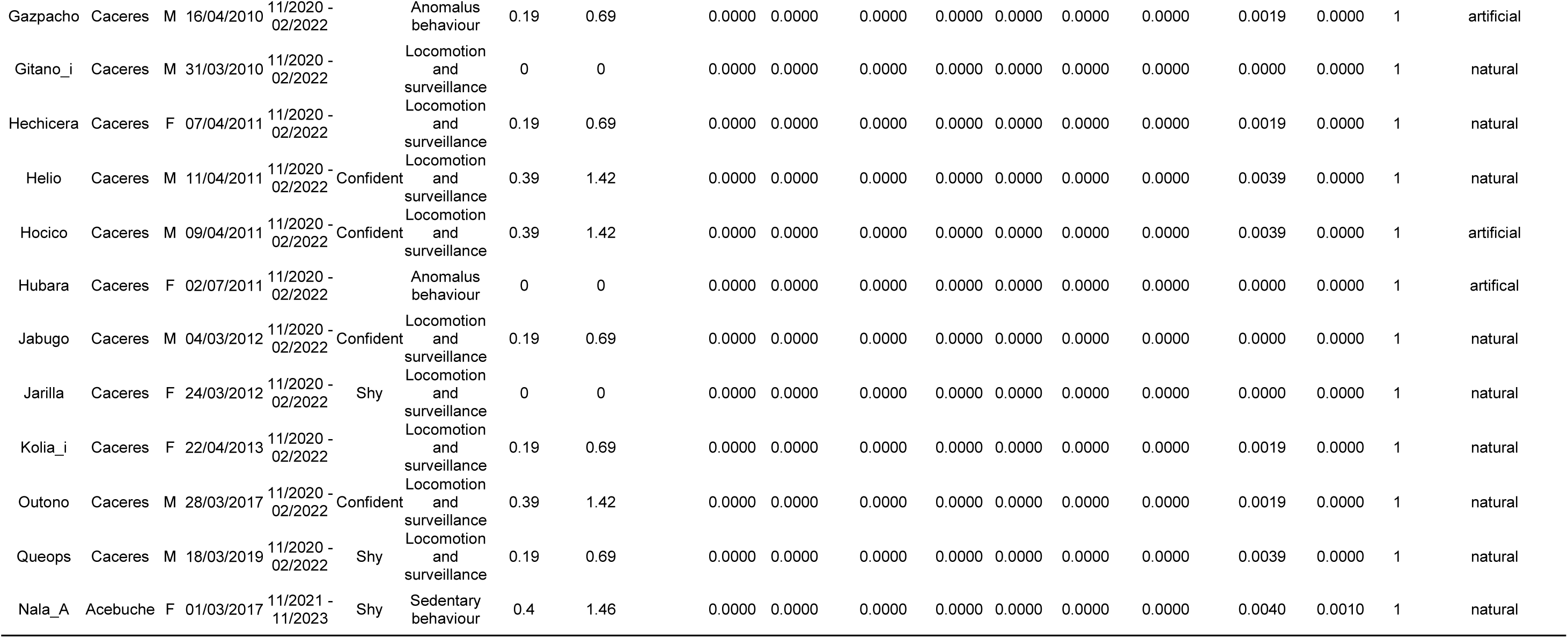

The individuals studied were preliminarily selected to cover “shy”, “confident” and “neutral” personalities according to the daily experience of the keeper staff (see Table 1). Some of the individuals were subjected to individualized drug treatments to deal with diseases and health disorders (Table 1, and see additional data in Supplementary Information). To include the variable “medication” in our study, we categorized and grouped drugs treatments as follows: (i) biotics (antibiotics and probiotics), (ii) analgesics (anti-inflammatories and painkillers), (iii) reproductive (to treat labor, lactation and chemical castration), (iii) drugs against renal insufficiency, (iv), gastroprotective (against nausea and heartburn), (v) nervous system (antidepressants, anxiolytics and antipsychotics), (vi) constipation, (vii) antiparasitic (against eukarya parasites), and (viii) vaccines (against viruses).

### 2.2. Behavior characterization

We initially selected 525,016 events of observation. From the initial pool, we deleted observations that both showed inter-individual interactions (Supplementary Fig. 1) and those events annotated as (i) not identified, (ii) out of sight, and (iii) out of camera range. The final curated behavioral dataset retained 430,917 events, equally distributed in Granadilla (211,741 events) and Acebuche (219,176). These observations were grouped according to the eight highest hierarchical levels previously defined in the standardized official ethogram as follows: (i) social, (ii) locomotion and surveillance, (iii) anomalous behavior, (iv) maintenance, (v) breeding season, (vi) feed, (vii) sedentary behavior, (viii) predatory behavior (inner crown, Supplementary Fig. 1). The behavioral observations were converted into frequencies, and a similarity matrix was created with vegdist function (Vegan package 2.6-6.1, Oksanen 2020), using the Euclidian method. A cluster analysis was run with the Nbclust function (from Nbclust package version 3.0.1, Charrad et al., 2014) and the ward.2 method, using all available indices and selecting the number of pre-defined clusters with the highest statistical support. A principal component analysis (PCA) (rda function, Vegan package 2.6-6.1) and a dendrogram (hclust function, package stats version 3.6.2) were additionally run to visualize the behavioral grouping. Finally, we ended with three behavioral categories, i.e., (i) anomalous, (ii) sedentary, and (iii) locomotion and surveillance. Additionally, Nbclust was applied for clustering lynxes according to medication treatments. Behavioral groups were compared using the variables sex, breading station, keeper cluster, drug treatment and upbringing, respectively, with Fisher and χ² tests (fisher.test and chisq.test functions, stats package version 3.6.2) (Agrasiti, 1990; Hope, 1968) and Cramer’s V (cramersV function lrs package; version 0.5.2) (Navarro, 2015). All statistical analyses were carried out in R (version 4.4.2).

Several months after the study ended, three individuals were translocated between sites (i.e., Nala from Acebuche to Granadilla, and Kolia and Gitano from Granadilla to Acebuche). We further analyzed potential changes in behaviour for such individuals just a few weeks before and several weeks after the translocation (Table 1).

### 2.3. DNA analyses

Fresh fecal samples were aseptically collected from the fenced areas twice per week, and were stored at -20 °C until further analysis. Samples were thawed and merged for biweekly periods after sub-sampling three different inner parts of the feces in a single composite sample (c., 1 gr wet weight). Subamples were placed in 2 mL cap tubes with 0.75 mL of lysis buffer (40 mM EDTA, 50 mM Tris pH 8.3, 0.75M sucrose) and c.a. 200 mg were extracted with the QIAamp DNA Stool Mini kit (QIAGEN, Hilden, Germany), and further analyzed by multiplexing PCR amplification and Illumina MiSeq sequencing following the methods of the Research Technology Support Facility at the Michigan State University (https://rtsf.natsci.msu.edu/). The V4 variable region of the 16S rRNA gene was amplified with primers F515 (5′-GTGCCAGCMGCCGCGGTAA-3′) and R806 (5′-GGACTACHVGGTWTCTAAT-3′) (Caporaso et al., 2011). A sequencing read length of 2x250 bp and paired end base pairs (bp) run was applied. Raw sequences were processed using the UPARSE pipeline (Edgar, 2013). Briefly, after merging of paired reads, filtering by an expected error of 0.25 and read length of 250 bp, ca. 69% of the original reads were retained (i.e., 18,039,709 reads). The reads were dereplicated and clustered into zero-radius operational taxonomic units (ZOTUs, identity 100%) excluding singletons. More than 98% of the merged sequence pool was mapped back into ZOTUs, and taxonomically assigned with SILVA_138.1 (Quast et al., 2013). Chloroplast, mitochondria, Eukarya, and unclassified reads were excluded for downstream analyses (< 0.03% of the reads mapped to ZOTUs). A total of 3,229 prokaryote ZOTUs were finally obtained. Only samples with at least 3,000 reads were retained, and the complete curated gut microbiota dataset ended with 647 samples. Laboratory blank control showed two orders of magnitude lower number of sequence-reads per sample than faeces samples, and an unrelated taxonomic profile (data not shown). The whole gene sequence dataset is available in the NCBI Sequence Read Archive through BioProject record ID PRJNA1481532

### 2.4. Statistical analyses for the microbial community and gut bacterial biomarkers

Microbial community parameters and multivariate statistics were calculated with the vegan package version 2.6-6.1 (Oksanen 2020). For distance-based community analyses, robust Aitchison metric (robust.aitchison) (Martino 2019) was calculated with vegdist function. The ordination representation was based on metric multidimensional scaling (cmdscale function). The beta-diversity, as distance to centroid for the different groups of samples, was calculated with betadisper and permutest functions. The permutational multivariate analysis of variance using distance matrices (adonis2 function) was carried out based on 1000 permutations to test for differences between groups of samples belonging to different factor categories, using the value of the statistic to evaluate the test performance since the p-value was found to be highly dependent on the number of permutations in most of the cases.

We used the vegdist matrix to evaluate differences in distance values (Welch’s t-test) between intra-group and inter-group comparisons for both behavior categories and breeding stations. We first conducted an exploratory analysis using descriptive statistics and boxplots, Levene’s test, and evaluating normality through Shapiro–Wilk tests and Q–Q plots, respectively. Effect sizes were computed using Cohen’s d with bootstrapped confidence intervals. We applied shapiro_test, t_test, and cohens_d functions from rstatix package (v. 0.7.3).

Richness and Shannon-Weaver index were calculated with the diversity and rrarefy functions (vegan package). All analyses were performed with a sampling depth of 3000 sequence reads per sample, and with the exception of richness, all were replicated hundreds of times using the replicate function from WRS2 package (v.1.1-6). Hypothesis contrast tests on the variance of the diversity indexes and Welch One-Way ANOVA test (tests for equal means in a one-way design without assuming equal variance) were performed using the function welch_anova_test with the rstatix package version 0.7.2. When needed, tests based on robust statistics methods were carried out, i.e. a one-way ANOVA on trimmed means using the t1way function (package WRS2, version 1.1-7), with 0.1 trims and 100 bootstraps. Corresponding post-hoc tests were run using the lincon function of the WRS2 package.

Differential abundance analysis suitable for compositional data was carried out using the package Analysis of Compositions of Microbiomes with Bias Correction (ancombc2 function within the package ANCOM-BC2 version 2.6.0) (Lin and Peddada, 2020). Previously, we created a TreeSummarisedExperiment object that contained a count table and categorical variables of the full metadata (TreeSummarisedExperiment package version 2.12.0) (Huang R, 2021). For the analysis, the breeding station was marked as a fix factor. Taxa <20% of occurrence and <0.01% of average abundance were deleted for the analysis. Figures were drawn with ggplot2 package version 3.5.2 (Wickham 2016).

We defined a taxonomic bacterial gut index based on the statistically significant differences in relative abundances found among the three behavioral categories according to both the ANCOM-BC2 analysis and the premise of at least 70% of occurrence. The best indicator taxa for each comparison was selected after Receiver Operating Characteristic (ROC) curves and calculating area under the curve (AUC) values, using the pROC package version 1.18.5 (Robin et al. 2011) (roc function). We tested each significant taxon individually and also in combination as follows (Taxon X + Taxon Y + 1e-6)/ (Taxon J + 1e-6). Optimal classification thresholds were determined using the Youden index (sensitivity + specificity − 1), which identifies the cutoff point that maximizes the combined performance of sensitivity and specificity. Data were shown in boxplots, winsorized at 10%. Hypothesis tests on the variance of the biomarkers were carried out as for the diversity indices, but using a 0.2 trim.

### 2.5 Structural equations model

To evaluate the direct and indirect effects of the breeding station on behavioural outcomes, we specified a Bayesian mediation model using the blavaan package in R (version 0.6-20, fuction bsem). The model included both the gut microbiota as intermediator expressed by the two components of the MDS analysis (V1 and V2), each predicted by the breeding station variable, and the three behavioral outcomes (anomalous, sedentary, and locomotion and surveillance, respectively), each regressed simultaneously on the intermediators and the predictor. Covariance between the mediators was freely estimated. For each behavioral outcome, we defined the specific indirect effects via V1 and V2, the combined indirect effect, and the total effect as the sum of direct and indirect components. Prior to estimation, all continuous variables were standardized to improve convergence and comparability of effects. The model was fitted using Bayesian structural equation modelling (BSEM) with blavaan, explicitly relying on Stan for posterior computation. Four Markov chains were run with 1,000 adaptation iterations, 2,000 burn-in iterations, and 5,000 post–burn-in samples per chain, providing 20,000 posterior draws for inference. Convergence was assessed through standard diagnostics.

### 2.6 Functional analyses

To approach functional predictions of the microbial assemblages, we used PICRUSt2 (standalone version) (Douglas et al., 2020). A 0.2 cutoff Nearest Sequenced Taxon Index (NSTI) value was chosen to increase prediction accuracy (Djemiel et al., 2022) after an assessment of the PICRUSt2 coverage in the original communities following different NSTI cutoff values. Briefly, we determined the degree of association of the original dataset and different subsets (generated at different NSTI cutoffs) by Mantel tests (function mantel in vegan R package Spearman method). Then, we evaluated the number of zOTUs retained at different NSTI cutoffs, and the percentage of abundance that was covered in the original communities (Fig supplementary 2). The functional analyses focused on predicted pathways abundances (folder output pathways-out). ANCOM-BC2 function for differential abundance analysis was used with breeding station as fixed factors and a 20% occurrence filter. We only selected those pathways showing a log fold change greater than 0.5 in more than one cluster comparison.

## RESULTS

### Behavioral traits and environmental effects

Overall, the behavioural dataset supported a strong three-cluster structure for all the Iberian lynx individuals analysed. After testing up to 30 indices available in NbClust, the strongest statistical result was obtained for three behavioural clusters supported by 10 indices (KL, CH, Hartigan, TrCovW, TraceW, Duda, PseudoT², Ball, Tau, SDindex), (Fig. 1, panel A), Each cluster was further characterized by mean behavioural frequencies and shown in radar charts (Fig. 1, panel B). Individuals in cluster 1 (yellow color in Fig 1) showed the highest frequencies for anomalous behaviour (>0.06), intermediate levels of locomotion and surveillance (0.20–0.32), and high frequencies of sedentary behaviour (>0.33). Individuals in cluster 2 (green color) were characterised by the highest sedentary behaviour frequencies (>0.44), intermediate locomotion and surveillance values (0.18–0.26), and low anomalous behaviour frequencies (<0.05). Finally, individuals in cluster 3 (purple color) showed low anomalous behaviour frequencies (<0.05), together with relatively balanced values for both sedentary behaviour (0.33–0.43) and locomotion and surveillance (0.32–0.39). Principal component analysis (PCA) provided a complementary ordination of the behavioural variation (Fig. 1, panel C), with the first two axes explaining 91.2% of the total variance (PCA1 = 47.8%; PCA2 = 43.3%). The main loading vectors in the biplot corresponded to three behavioural dimensions, i.e., anomalous, sedentary, and locomotion and surveillance behaviours, respectively. The spatial distribution of individuals across the PCA confirmed the separation of the clusters and allowed their behavioural allocation. Accordingly, cluster 1 was associated with the anomalous behaviour axis (AB), cluster 2 with the sedentary behaviour axis (SB), and cluster 3 with the locomotion and surveillance axis (LS), in agreement with the radar plots. Additional evidence from translocated individuals showed both stability and change in behavioural classification. Nala and Gitano maintained their original behavioural clusters (SB and LS, respectively) after translocation between breeding stations. In contrast, Kolia shifted from the LS cluster to the AB cluster following translocation.

**Figure 1.**
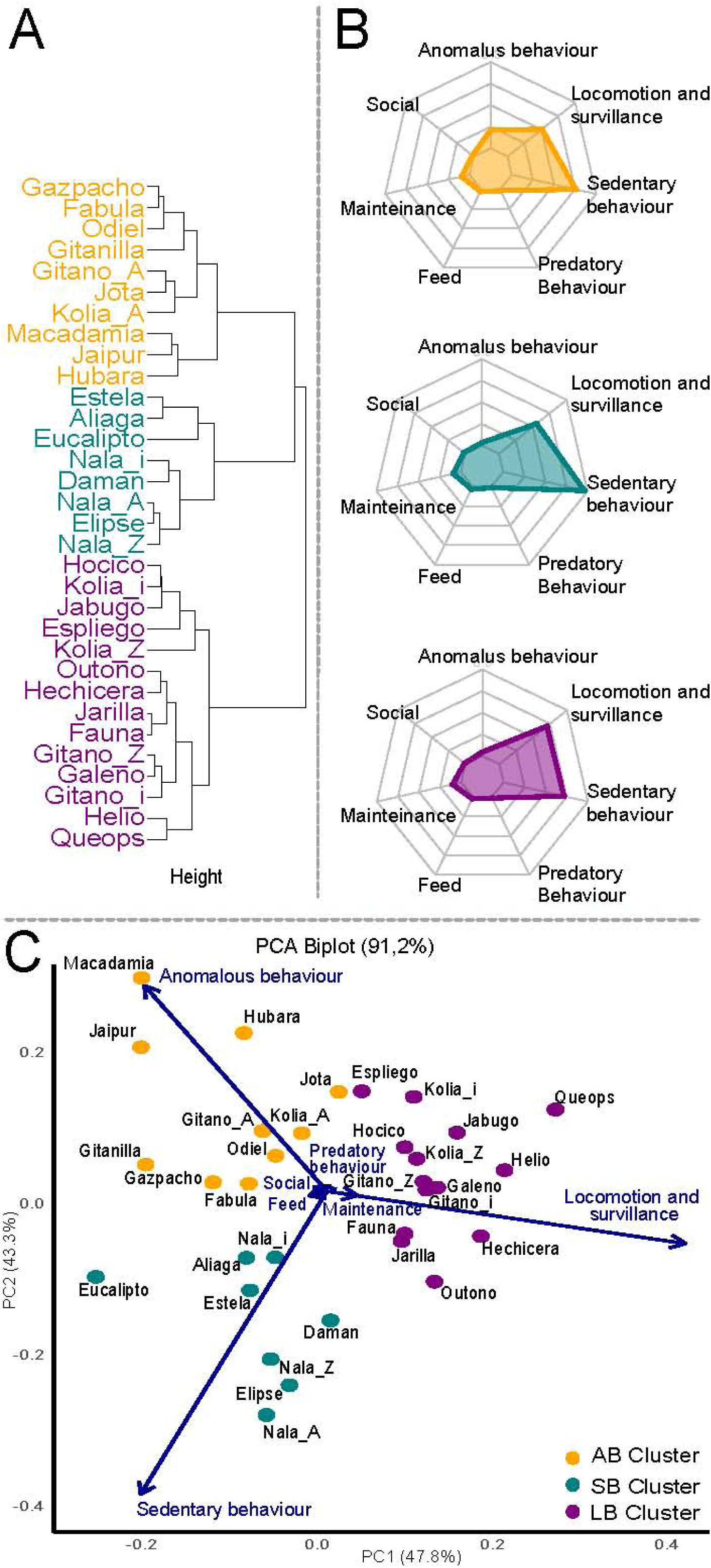
Clustering of Iberian lynx individuals based on behavioural frequencies. (A) Euclidean distance-based hierarchical clustering dendrogram. Individuals are coloured according to the three behavioural clusters identified. (B) Radar plots illustrating the mean frequency for each behavioural category coloured according to the three clusters identified. (C) Principal component analysis (PCA) biplot based on individual behavioural frequencies. Points represent individual observations and are coloured according to behavioural cluster membership, whereas blue vectors indicate the contribution and direction of each behavioural variable. Percentages of variance explained by the first two principal components are indicated on the axes. A few individuals were translocated between breeding stations in a secondary experiment and are labelled as “_I” (main experiment), “_A” (Acebuche, secondary experiment), and “_Z” (Zarza de Granadilla, secondary experiment).

Individuals within the breeding program are subjected to veterinary treatments and, therefore, medication intake need to be considered as a potential explanatory variable. Overall, medication was low across individuals and treatment categories (Supplementary Fig. 3B), and only a few individuals showed comparatively high medication ratios, i.e., Odiel (associated with nervous system treatments), and Jota, Fabula, and Espliego (related to renal insufficiency treatments) had intake frequencies close to 0.2. Exceptionally, Eucalipto showed an intake frequency of 0.87 for renal insufficiency illness. A cluster analysis based on medication intake provided strong support for a two-cluster structure, as agreed by eight NbClust indices (KL, DB, ptbiserial, McClain, gamma, Gplus, Dunn, and SDindex) where Eucalipto split from all remaining counterparts.

We further assessed whether behavioural clusters were associated with additional explanatory variables, including sex, breeding station, keeper-based classification, medication cluster, and upbringing. This analysis was used to evaluate the robustness of the behavioural clustering and to identify potential interactions beyond the primary relationship with gut microbiota. Among the variables tested, statistically significant associations were detected for breeding station (p_F_-value = 0.00, V = 0.674) and upbringing (p_F_-value = 0.01, V = 0.56), both showing strong effect sizes. Conversely, no significant associations were found for sex (p_F_-value = 0.40, V = 0.31) or medication ratios (p_F_-value = 0.24, V = 0.36). The association with keeper-based classification was marginal (p_F_-value = 0.05, V = 0.57), suggesting partial agreement between the behavioural clusters inferred in this study and the daily observations made by keeper staff. Overall, the high effect size and Cramér’s V values detected for both breeding station and upbringing (>0.55) supported a strong relationship between these explanatory variables and behavioural clustering (Supplementary Fig. 3A).

We also compared climatic variables and vegetation cover between the two breeding stations (Supplementary Fig. 4). Overall, environmental conditions were broadly similar during the study period, with only a limited number of significant differences. Dewpoint temperature differed between stations across all seasons (13.25 ≥ t ≥ 3.23; 10.46 ≥ df ≥ 2.68; p-value ≤ 0.02; Supplementary Fig. 4, d2m panel), and the eastward wind component differed in spring, summer, and autumn (7.73 ≥ t ≥ 3.00; 9.10 ≥ df ≥ 6.19; p-value ≤ 0.02; u10 panel). In both cases, Acebuche showed higher values than Granadilla. Additional seasonal differences were detected in winter. Temperature was lower (t = 6.34; df = 9.23; p < 0.01), whereas evaporation was higher (t = -4.28; df = 9.56; p < 0.01) in Granadilla The northward wind component also reached the highest values in Acebuche during summer. Finally, vegetation cover was higher in Granadilla than in Acebuche; however, no statistical test was applied because both vegetation-cover variables had zero variance.

### Gut microbial communities

MDS ordination based on gut microbiota community composition showed only limited separation among Iberian lynxes assigned to the three behavioural clusters (Fig. 2A; adonis test R^2^ = 0.011; p = 0.001). The three groups largely overlapped in ordination space, but SB and LS showed a slight separation from each other. Community distances within each behavioural cluster were slightly lower than distances between clusters, indicating greater similarity among samples from the same behavioural group. This pattern was supported by significant intra- versus inter-group comparisons for AB (Welch’s t-test: t = 30.6, df = 92,455, p < 0.001; negligible effect size = 0.170), SB (Welch’s t-test: t = 4.74, df = 30,316, p = 2.2 × 10^-6^; negligible effect size = 0.0369), and LS (Wilcoxon test: statistic = 429,883,793, df_1_ = 1, df_2_ = 80,708, p = 0; small effect size = 0.138). Together, these results suggest modest but consistent microbiota differences among behavioural clusters.

**Fig 2.**
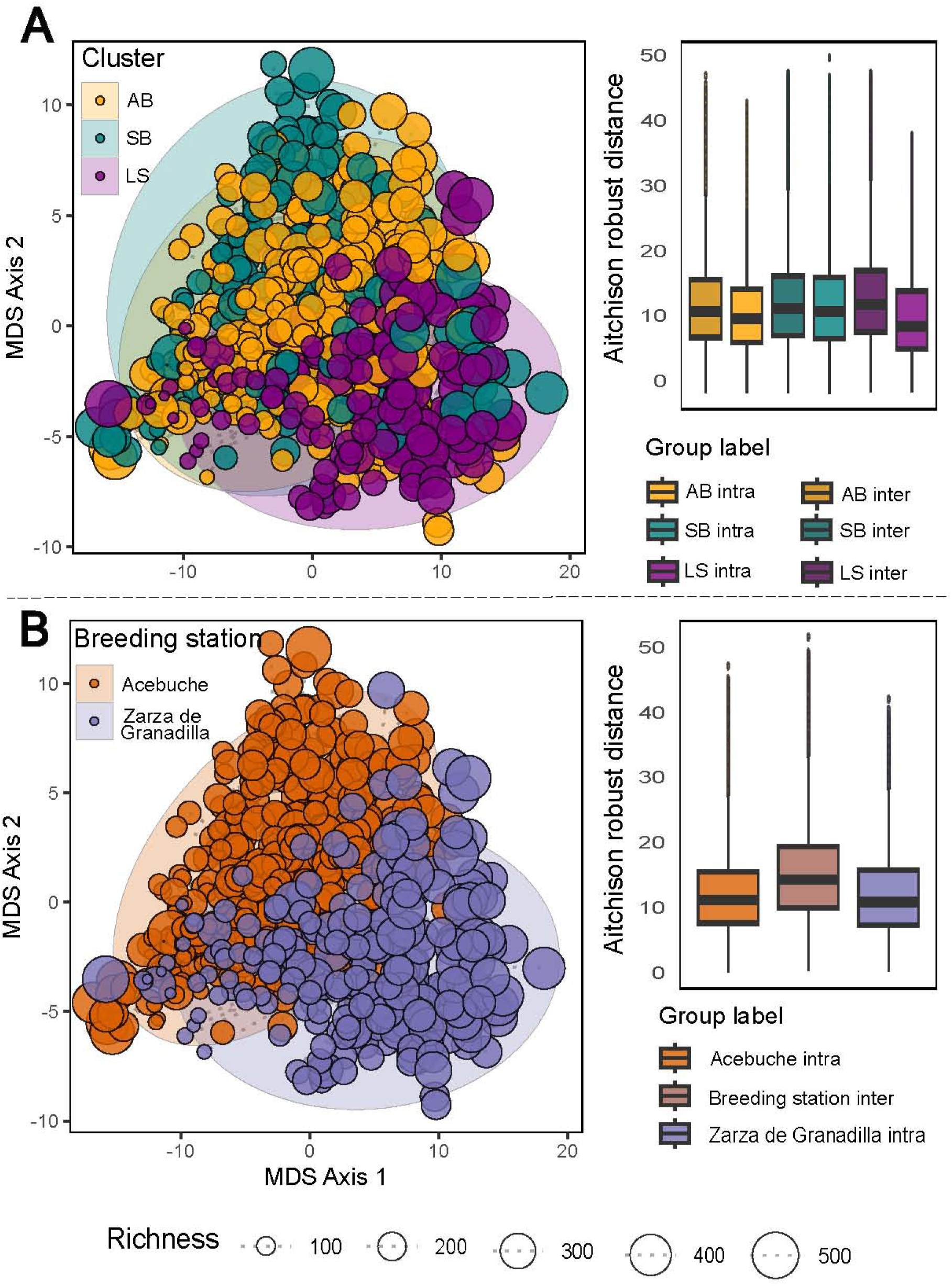
Gut microbiota multidimensional scaling ordination (MDS) based on robust Aitchison distances, highlighted by behavioural profiles (panel A) and breeding stations (panel B). Right panels show inter- and intra-comparisons of distances.

A similar pattern was observed when samples were grouped by breeding station (Fig. 2B). MDS ordination showed weak separation between Acebuche and Granadilla (adonis R^2^ = 0.038, p = 0.001). Intra-station distances were slightly lower than inter-station distances, indicating that samples from the same breeding station were more similar to each other than to samples from the other station for both Acebuche (Welch’s t-tests t = -96.0, df = 179,209, p = 0; small effect size = 0.452) and Granadilla (t = 66.8, df = 49,646, p = 0; small effect size = 0.440) although the magnitude of the separation were moderate to small. Conversely, upbringing practices poorly explained variation in gut microbiota composition. The multivariate analysis showed negligible group separation (ANOSIM, R = 0.02, p = 0.002). Thus, the effect of upbringing on microbial gut community structure was weaker than the effect of breeding station.

The Bayesian structural equation model (Fig. 3) showed satisfactory fit and convergence (PPP = 0.508, marginal log-likelihood = 3884.71, R^ ≈ 1.00). Shifting from Acebuche to Granadilla was associated with an increase in V1 (β = 0.466; 95% CI: 0.826 to 1.105) and a decrease in V2 (β = -0.513; 95% CI: -1.200 to - 0.926). Gut-microbiota mediation varied by behaviour. For anomalous behaviour, indirect effects were weak and not significantly different from zero. Instead, the direct effect was stronger and negative, resulting in an overall negative total effect (standardised β = -0.318; 95% CI: -0.810 to -0.507). For sedentary behaviour, V1 showed only a weak and non-significant association, whereas V2 had a stronger contribution to the model, producing a negative total effect (standardised β = -0.258; 95% CI: -0.537 to -0.384). The strongest mediation pattern was observed for locomotion and surveillance behaviour. In this case, microbiota components contributed positively, leading to a positive total effect (standardised β = 0.449; 95% CI: 0.788 to 1.074). Thus, breeding station influences behaviour through both direct environmental effects and indirect microbiota-mediated pathways, with the latter most evident for locomotion and surveillance.

**Figure 3.**
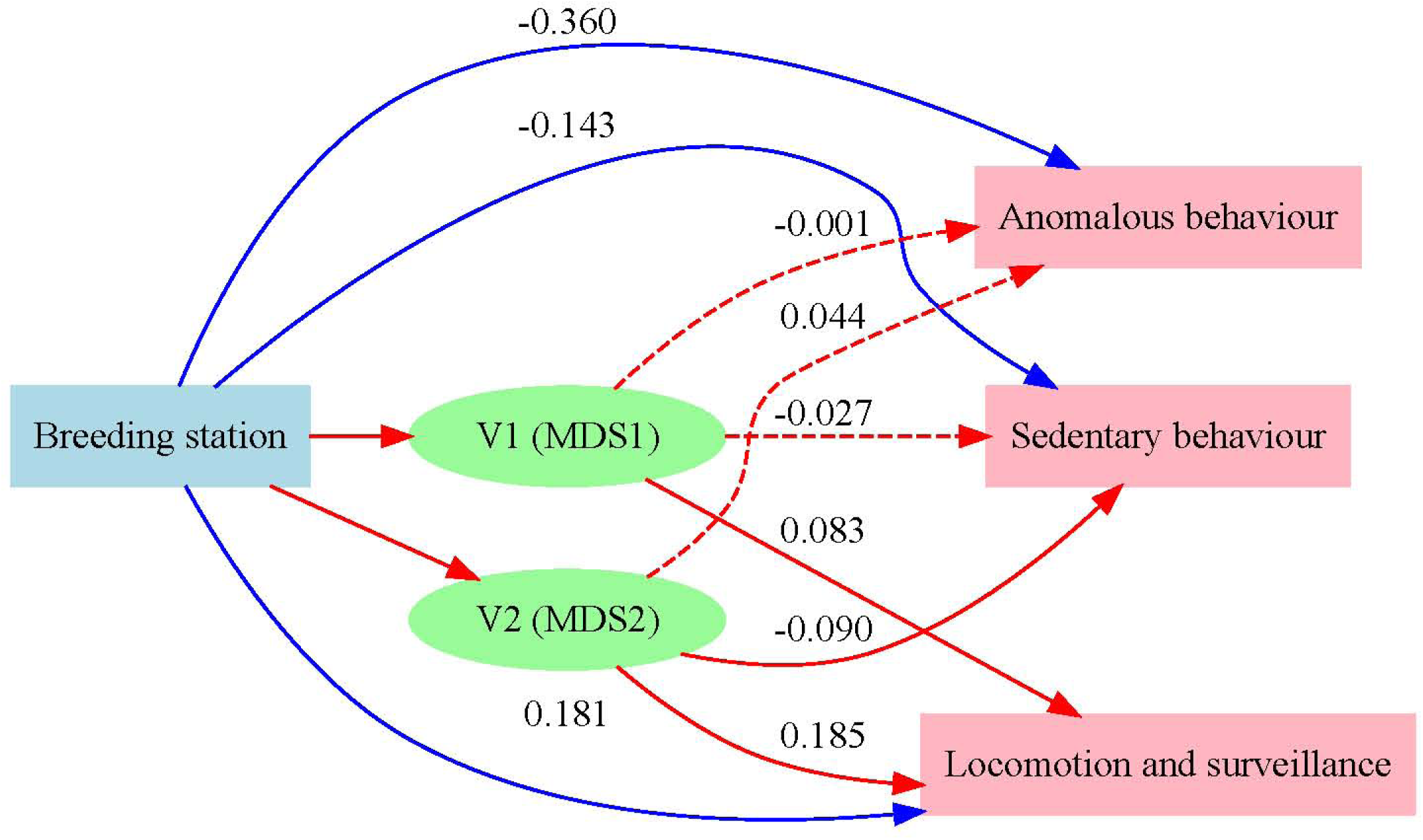
Bayesian structural equation model showing relationships bewteen breeding environment, gut microbiome and behavioural phenotypes, respectively. Blue arrows indicate direct effects of breeding station (blue box) on behavioural traits (red boxes), whereas red arrows represent indirect effects mediated through gut microbiome composition (green circles). Solid arrows mean statistically supported relationships, whereas dashed arrows indicate non-significant paths.

Alpha ana Beta-diversity analyses revealed significant differences among behavioural clusters for betadiper (distance to centroid), species richness, and ecological diversity measured with the Shannon index (Supplementary Fig. 5A). For betadisper, robust one-way ANOVA showed significant differences among clusters (t1way test: F = 5.33, df_1_ = 2, df_2_ = 274.05, p = 0.005), although the effect size was small (effect size = 0.20). Post hoc comparisons based on trimmed means indicated significant differences between AB and LS (psihat = - 1.29, 99% CI [-2.49, -0.09], p = 0.005) and between SB and LS (psihat = -1.22, 99% CI [-2.58, 0.14], p = 0.018), whereas AB and SB did not differ significantly (psihat = -0.07, 99% CI [-1.20, 1.06], p = 0.854). Species richness showed the same general pattern as betadisper. Differences among clusters were significant (t1way test: F = 5.40, df_1_ = 2, df_2_ = 266.62, p = 0.005), with a small effect size (effect size = 0.22). Post hoc tests showed significant differences for AB versus LS (psihat = -10.66, 99% CI [-20.97, -0.34], p = 0.006) and SB versus LS (psihat = -11.35, 99% CI [-22.25, -0.45], p = 0.006), but not for AB versus SB (psihat = 0.69, 99% CI [-7.41, 8.79], p = 0.804). For the Shannon index, robust one-way ANOVA also detected significant differences among clusters (t1way test: F = 4.54, df_1_ = 2, df_2_ = 278.46, p = 0.011), again with a small effect size (effect size = 0.19). However, only the comparison between SB and LS was significant (psihat = -0.22, 99% CI [-0.44, -0.01], p = 0.009), whereas AB versus SB (psihat = 0.10, 99% CI [-0.08, 0.27], p = 0.116) and AB versus LS (psihat = -0.13, 99% CI [-0.33, 0.01], p = 0.116) were not significant. Overall, the LS cluster consistently had the highest values for all three diversity indices, indicating greater internal variability, higher species richness, and greater ecological diversity. This pattern reinforces the separation of LS from AB and SB seen in behavioural clustering. When samples were grouped by breeding station (Supplementary Fig. 5B), Granadilla showed higher microbial diversity than Acebuche. All three indices differed significantly between stations. Betadisper showed a moderate effect size (t1way test: t = 28.664, df_1_ = 1, df_2_ = 369.1, p = 0; effect size = 0.31), as did species richness (t1way test: t = 24.298, df_1_ = 1, df_2_ = 363.07, p = 0; effect size = 0.30). The Shannon index was also significantly different between stations, although the effect size was small (t1way test: t = 15.583, df_1_ = 1, df_2_ = 392.82, p = 9 × 10^-5^; effect size = 0.21).

### Compositional differences in bacterial taxa among behavioural groups and breeding stations

Compositional differences in bacterial taxa were observed for the different behaviour clusters (Fig. 4). ZOTUS related to *Acinetobacter, Peptostreptococcus, Peptoestreptococcaceae unclassified, Anaerovoracaceae;S5-A14a, Butyricicoccus, Lachnospiraceae NK4A136 group, Lachnoclostridium, Blautia, Desulfovibrio, Parabacteroides,* and *Bacteroides* were significantly more abundant in the AB cluster than in the SB cluster. More, *Blautia* and *Lachnospiraceae unclassified* were more abundant in AB than in LS cluster. *Negativacillus, Faecalibacterium*, and *Lachnospiraceae unclassified* were more abundant in the AB cluster than in the other groups, whereas *Lachnospiraceae*;*GCA-900066575* was less abundant than in the other groups. *Fusobacterium* is more abundant in SB than in LS, whereas *Eggerthellaceae; unclass* and *Collinsella* are more abundant in SB than in AB. *Lachnoclostridium, [Ruminococcus] gnavus group, [Ruminococcus] gauvreauii group, Allobaculum, Alloprevotella*, and *Bacteroides* are more abundant in SB than in the other groups, whereas *Fusobacterium, UCG-005, Lachnospiraceae unclass, Clostridiaceae unclass, Clostridia UCG-014 unclass, Parabacteroides, Prevotella_7*, and *Alloprevotella* are lower in this cluster.

**Figure 4.**
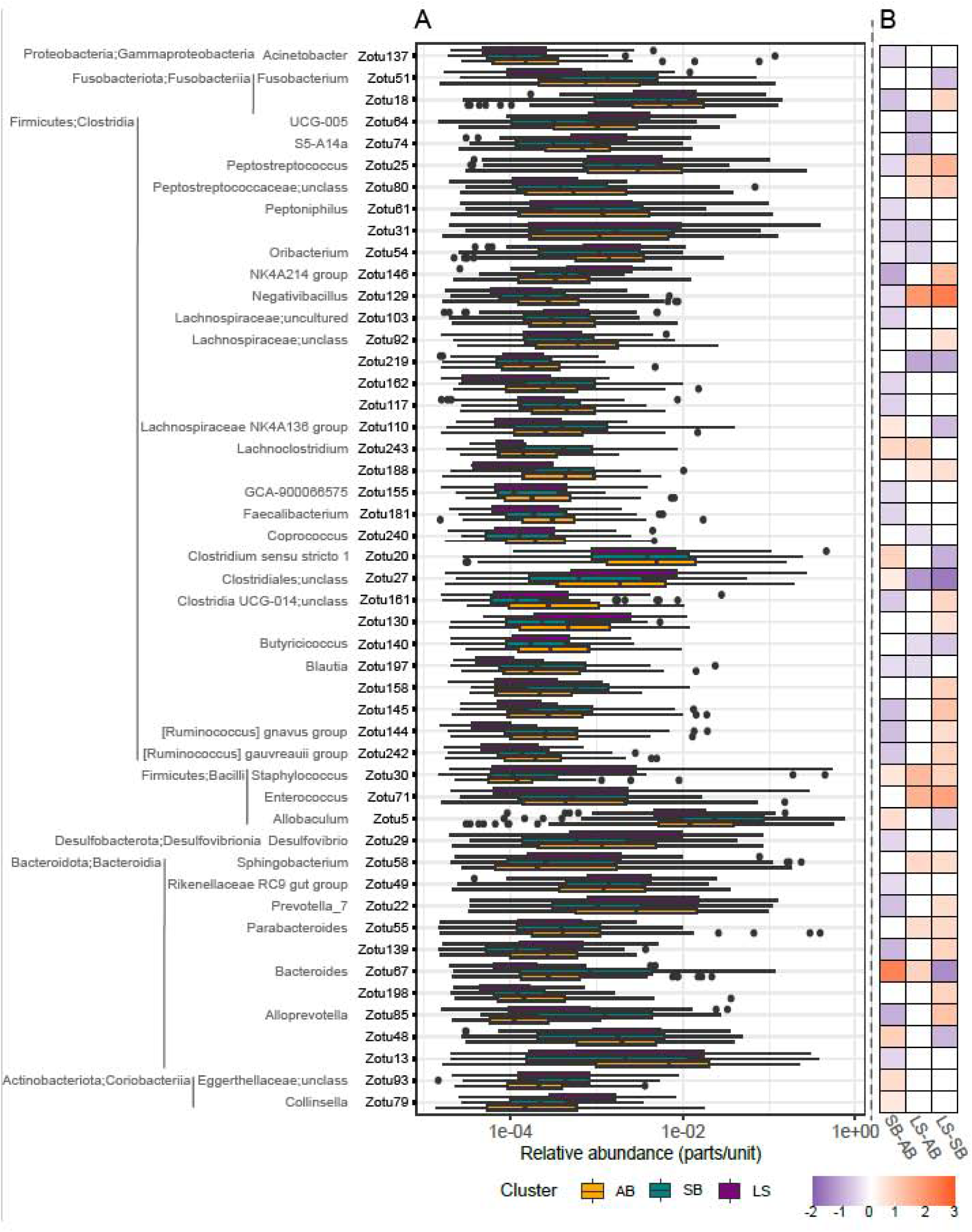
ANCOM-BC2 data for relative abundances of bacterial ZOTUs showing significant differences among behavioural clusters (panel A). Panel B shows a heatmap summarising ANCOM-BC2 results analysis where columns are pairwise comparisons between behavioural clusters, and rows show statistically significant associations. Cell colour indicates the direction and magnitude of the estimated log-fold change, with purple indicating lower abundance and orange indicating higher abundance.

**Figure 5.**
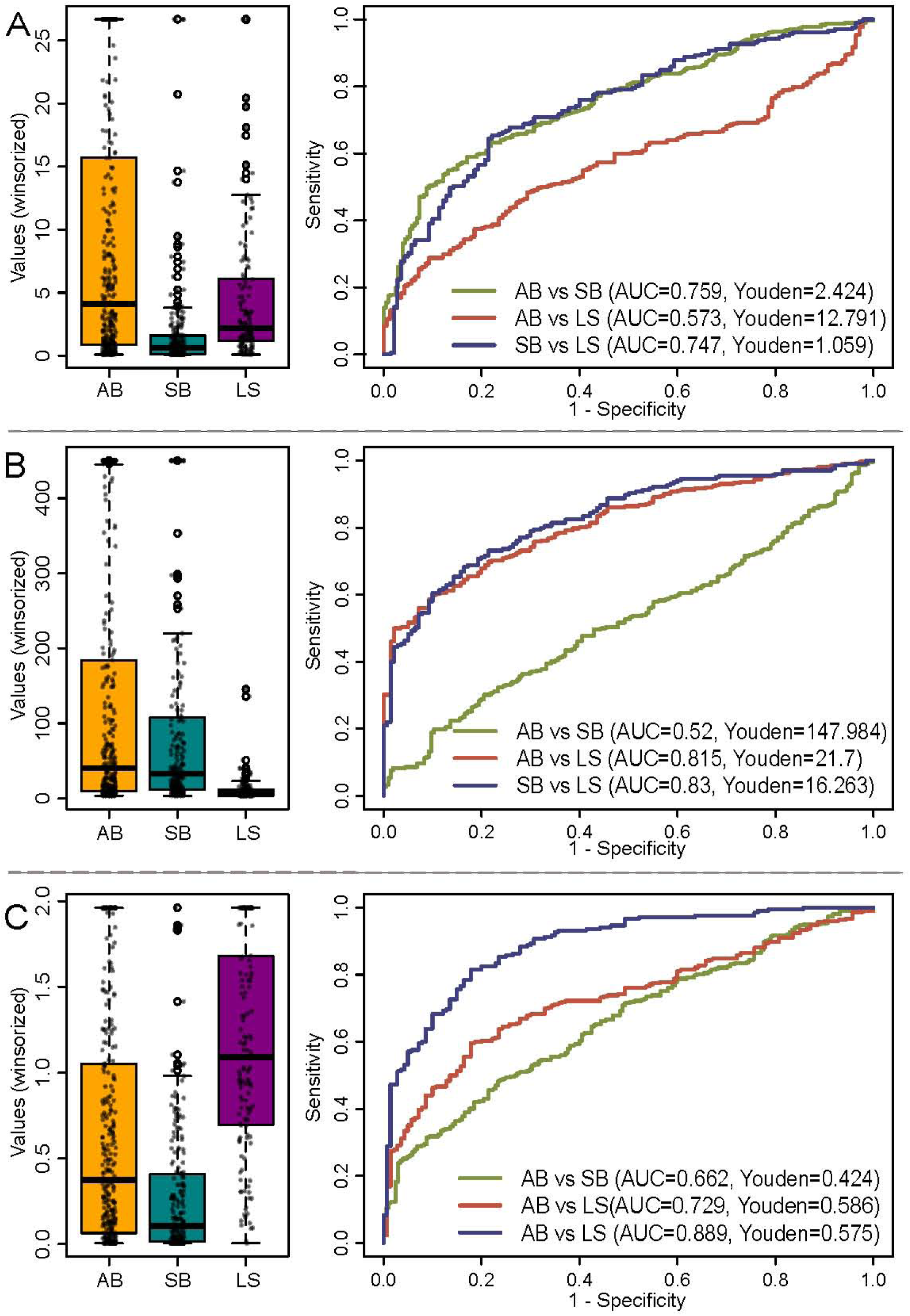
Taxonomic log-ratios discriminating behavioural clusters. Left panels show the distribution of winsorized taxonomic log-ratios across behavioural clusters. Right panels present Receiver Operating Characteristic (ROC) curves evaluating the discriminatory performance of each ratio for pairwise comparisons among anomalous behaviour (AB), sedentary behaviour (SB) and locomotion-and-surveillance (LS) clusters. Area under the curve (AUC) and the corresponding Youden index are reported for each comparison. (A) (Erysipelatoclostridiaceae; UCG-004 + Oscillospiraceae; UCG-005 + 1 × 10⁻⁶)/(Coriobacteriaceae; Collinsella + 1 × 10⁻⁶). (B) (Erysipelotrichaceae; Allobaculum + Lachnospiraceae; Blautia + 1 × 10⁻⁶)/(Coriobacteriaceae; Collinsella + 1 × 10⁻⁶). (C) (Erysipelotrichaceae; Solobacterium + Oscillospiraceae; UCG-005 + 1 × 10⁻⁶)/(Erysipelotrichaceae; Allobaculum + 1 × 10⁻⁶).

*Lachnospiraceae;unclultured, Lachnospiraceae;unclass, Clostridium sensu stricto 1*, and *Alloprevotella* are more abundant in LS than in SB clusters*. Peptoniphilus, Oscillospiraceae;NK4A214 group, Coprococcus, Staphylococcus, Enterococcus, Sphingobacterium*, and *Rikenellaceae RC9 gut group* are more abundant in LS cluster than in the others, whereas *Oribacterium* and *Lachnospiraceae;unclass* are less abundant. From these, the most represented genera in the analysis (i.e., each with more than one Zotu) included Fusobacteria, Peptoniphilus, Lachnoclostridium, Blautia, Lachnospiraceae, Clostridia UCG-014; unclassified, Parabacteroides; unclassified, Alloprevotella, and Bacteroides.

We also obtained a list of different ZOTUs unevenly distributed across the breeding stations (Supplementary Fig. 6). A comparison of the two analyses showed common biomarkers shared between behavioural groups and breeding stations, namely: *Peptoniphilus, Anaerovoracaceae S5-A14a, Negativibacillus, UCG-005, NK4A214 group, Lachnoclostridium, Blautia, [Ruminococcus] gnavus group, [Ruminococcus] gauvreauii group, Clostridium sensu stricto, Staphylococcus, Enterococcus, Allobaculum, Desulfovibrio, Parabacteroides, Rikenellaceae RC9 gut group, Prevotella 7, Alloprevotella, Bacteroides* and *Collinsella* (uncultured and unclassified genera were excluded).

### Functional analysis

We searched for different distribution of metabolic pathways along behaviour clusters by means of PICRUSt2 inferred functional data. We repeated the same differential abundance analysis strategy as done with the taxa. We obtained a list of pathways with significantly different abundances among the behaviour clusters (Supplementary Table 1). We can group the 14 pathways in 5 groups: (I) degradation of amino acids and phosphate compounds, (II) degradation of aromatic compounds, (III) carbohydrate metabolism, (IV) Lipids/lipopolysaccharide biosynthesis, (V) Biosynthesis of wall compounds. The pathways that are more abundant in SB and LS than in AB are: L-leucine degradation I (group I) and 3-phenylpropanoate degradation (group II). In LS, S-methyl-5-thio-α-D-ribose 1-phosphate degradation, L-methionine salvage cycle III (booth group I), glycogen degradation II, sucrose biosynthesis III, sucrose biosynthesis I (from photosynthesis) (all within group III of pathways), superpathway of geranylgeranyldiphosphate biosynthesis I (via mevalonate), mevalonate pathway I (booth group IV), peptidoglycan biosynthesis II (staphylococci), and peptidoglycan biosynthesis V (β-lactam resistance) (booth group V), are more abundant in LS than in AB and SB. In SB and AB, L-tryptophan degradation XII (Geobacillus) (group I) and 2-aminophenol degradation (group II) are more abundant than in LS. The superpathway of lipopolysaccharide biosynthesis is more abundant (group III) in AB than in the other clusters.

### Bacterial gut biomarkers discerning between behaviour groups

We identified 23 genera that differed significantly among the three behavioural clusters (Supplementary Fig. 7). We then tested several genus combinations to identify the most informative biomarker ratios for behavioural classification (Fig. 6). For AB versus SB clusters : the ratio (*Erysipelatoclostridiaceae;UCG-004* + *Oscillospiraceae;UCG-005* + 1×10⁻⁶) / (*Coriobacteriaceae;Collinsella* + 1×10⁻⁶) achieved 75.9% accuracy. Values above 2.424 classified samples as AB, whereas lower values classified them as SB. In the case of AB versus LS clusters: the ratio (*Erysipelotrichaceae;Allobaculum* + *Lachnospiraceae;Blautia* + 1×10⁻⁶) / (*Coriobacteriaceae;Collinsella* + 1×10⁻⁶) reached 81.5% accuracy. Values above 21.7 indicated AB, whereas lower values indicated LS. For SB versus LS clusters: the ratio (*Erysipelotrichaceae;Solobacterium* + *Oscillospiraceae;UCG-005* + 1×10⁻⁶) / (*Erysipelotrichaceae;Allobaculum* + 1×10⁻⁶) achieved 88.9% accuracy. Values above 0.575 classified samples as LS, whereas lower values classified them as SB. Accuracy values and classification thresholds were also calculated for all pairwise behavioural-cluster comparisons using the three selected genus combinations. Alternative ratios are reported in Fig. 6, although they showed lower discriminatory performance than the combinations described above.

All selected biomarker ratios were supported by robust statistical tests. The first ratio showed strong differences among clusters (t1way test: t = 29.806, df_1_ = 2, df_2_ = 219.81, p = 0.000; effect size = 0.62, large), with significant pairwise differences between AB and SB (lincon test: psihat = 9.01, 99% CI [4.90, 13.12], p < 0.000). The second ratio also differed significantly among clusters (t1way test: t = 35.865, df_1_ = 2, df_2_ = 262.17, p ≤ 0.02059; effect size = 0.43, moderate), with significant differences between AB and LS (lincon test: psihat = 100.65, 99% CI [61.64, 139.66], p < 0.000). Finally, the third ratio showed strong differences among clusters (t1way test: t = 68.513, df_1_ = 2, df_2_ = 238.02, p = 0.000; effect size = 0.63, large), with significant differences between SB and LS (lincon test: psihat = -1.03, 99% CI [-1.26, -0.79], p < 0.000).

## DISCUSSION

Although growing evidence supports bidirectional interactions between the gut microbiome, behaviour, and environmental conditions, these components are generally investigated independently (Mayer et all; 2015). Most microbiome studies have been conducted under laboratory conditions or have focused on human health and neurological disorders (Cryan et al., 2019; Davidson et al., 2020; He et al., 2024), whereas studies integrating behavioural ecology and microbial ecology in wild or conservation dependent species remain scarce. Besides, environmental variation has been shown to influence gut microbial communities (Heni et al., 2023). While pairwise relationships between environment and behaviour, environment and microbiota, or microbiota and behaviour have been increasingly explored, studies simultaneously integrating all three components within a conservation framework remain virtually absent, particularly in endangered carnivores. In this study, we integrated behavioural ecology, microbial ecology, and environmental variables, providing a comprehensive framework for help to understand the mechanisms underlying individual behavioural variation. This integrative approach could be particularly relevant for conservation programmes, where understanding individual variability is essential to optimise management decisions and improve reintroduction success. Behaviour-focused studies have traditionally emphasized dispersal and home-range movements; however, they also highlight the importance of incorporating personality traits to achieve a more comprehensive understanding of how animals interact with their environments. In this context, integrating personality classification with movement-tracking methodologies, may provide deeper insights into how individual perceive and respond to environmental conditions, ultimately offering a more holistic framework for understanding individual ecological strategies (Cisneros-Araujo, 2025).

Previous studies on Iberian lynx behaviour and dispersal had mainly described long-range movements associated with settlement search, home-range movements within established areas, and variability within these movement patterns (Rueda, 2011). Other work had emphasized the importance of individual traits and personality in exploratory and settlement movements (Wolf et al., 1996; Yiu et al., 2015). The identification of these two types of movements in wild environments, as well as their variability, is consistent with our findings on behavioural personality in captive individuals. Specifically, lynxes within the SB cluster showed shorter or less frequent movements, whereas individuals in the LS cluster tended to display longer-range or more frequent movements. Conversely, anomalous behaviours, or stereotypies, are well documented in the literature and may appear in captive animals as responses to stress. Importantly, these behaviours can persist after reintroduction into the wild, potentially reducing reintroduction success (Vaz et al., 2017; Fischer et al., 2019; Vickery et al., S 2003). Thus, clusters SB and LS appear to represent behavioural phenotypes that are more closely aligned with wild-type patterns, whereas the AB cluster reflects a less common profile under natural conditions due to its association with stereotypic behaviours. In this context, identifying lynxes that exhibit high frequencies of abnormal behaviours can support Iberian lynx reintroduction efforts by detecting individuals with a lower likelihood of successful release.

Previous studies in felids have reported behavioural differences between cubs reared with and without maternal care (Bouchet et al., 2022; Hampson et al., 2016), and research on the Iberian lynx has likewise documented differences in growth between artificially and naturally reared individuals (Yerga J., 2014). This is consistent with our findings, which reveal a positive association between the AB cluster and artificially reared individuals. However, our results also indicate a clear relationship between breeding centre and behavioural phenotype, i.e., individuals belonging to the AB cluster were more frequent in Acebuche, whereas the LS cluster was absent. This asymmetric distribution raises an important question: does it reflect differences in observation or a genuine environmental effect? Comparisons between the two stations indicated that Acebuche was characterized by stronger winds and lower vegetation cover than Granadilla. In the wild, these environmental features are relevant because Iberian lynxes usually select areas with vegetation cover, including trees, shrubland, and grassland (Cisneros-Araujo, 2025). In addition, greater landscape heterogeneity has been associated with a broader range of behavioural phenotypes (Mortelliti A, 2020), while above-average wind speeds tend to modify normal behavioural patterns in Canada lynx in boreal forests (Studd EK, 2022). Enclosures with greater environmental heterogeneity tend to promote higher behavioural diversity and reduce the occurrence of abnormal behaviours, whereas less complex environments or adverse environmental conditions can alter activity patterns and space use (Fàbregas et al.,2012). Taken together, these observations suggest that the initial asymmetric distribution of behavioural clusters between breeding stations may be linked to environmental conditions rather than to methodological bias on taking behavioural data. In particular, stronger winds combined with reduced vegetation cover in Acebuche may limit shelter availability and landscape heterogeneity, potentially altering behavioural expression. By contrast, the greater vegetation cover and environmental heterogeneity in Granadilla may provide conditions more similar to natural Iberian lynx habitats, which could help explain the presence of LS individuals at this station.

The gut microbiome analysis revealed a clear association between microbial community composition and breeding station of origin. This pattern was expected, as previous studies have shown that geographic location shape gut microbiota structure across animal hosts (Wang et al.,2022; Xie et al., 2024). In our study, the breeding station therefore appears to represent an important environmental context influencing gut microbial variation. We also detected associations between gut microbiome composition and behavioural phenotype. This finding is consistent with a growing body of evidence linking gut microbial communities with neural, physiological, and behavioural conditions (Verma A., 2024). Although such associations must be interpreted cautiously, they suggest that microbiome variation may contribute to, or reflect, individual behavioural differences in Iberian lynxes. Several aspects of our experimental design reinforce the robustness of these interpretations and reduce the likelihood that the observed patterns arise from spurious or confounding effects. First, the large sample size provides substantial statistical power, increasing the reliability and reproducibility of the detected associations. This extensive dataset reduces the influence of stochastic variation and allows subtle but biologically meaningful patterns to emerge. Second, behavioural characterization was not based on isolated or short-term observations, nor reduced to single categorical or clinical-type measures. Instead, behaviour was quantified through continuous, standardized, and repeated observations over extended periods of time, capturing consistent behavioural tendencies across temporal scales. This longitudinal approach provides a more accurate representation of individual behavioural phenotypes and minimizes the impact of transient or situational responses. Third, the study design explicitly incorporates environmental heterogeneity by including individuals housed in distinct breeding centres and by accounting for temporal variation. This allows us to partially disentangle the effects of environmental context from intrinsic individual differences. In addition, the integration of multiple explanatory variables reduces the risk of oversimplified interpretations based on single-factor associations. Importantly, the convergence of results across behavioural, microbial, and environmental datasets further strengthens the validity of our findings. Rather than relying on a single line of evidence, the consistency of the observed patterns across independent analytical approaches supports the biological relevance of the associations. Taken together, these methodological strengths provide a robust framework that increases confidence in our results and supports the interpretation that the detected behavioural–microbiome relationships reflect genuine biological processes, rather than being solely driven by uncontrolled environmental variation or methodological bias.

Looking now to the three lynxes that were translocated between breeding stations: Nala, Kolia, and Gitano, for further explore the potential effects of environmental change on individual behaviour. Two individuals maintained their original behavioural clusters after translocation: Nala remained in the SB cluster, and Gitano remained in the LS cluster. In contrast, Kolia shifted from the LS cluster to the AB cluster after moving to a different environmental context. These cases provide a preliminary opportunity to evaluate whether behavioural classification is stable across environments or may change in response to environmental conditions. The persistence of Gitano within the LS cluster after translocation from Granadilla to Acebuche also suggests that the behavioural patterns detected in this study are unlikely to result from observer bias. Gitano retained the LS phenotype despite being moved to a station where no other lynx was classified as LS, supporting the consistency of the ethological sampling and behavioural clustering approach. Based on these observations, two non-mutually exclusive mechanisms may explain the individual responses observed after translocation. First, behavioural stability in individuals such as Gitano and Nala may reflect resilience of the gut microbiome and its associated influence on behaviour, even under changing environmental conditions. Second, the shift observed in Kolia suggests that environmental change, potentially together with microbiome restructuring, may contribute to changes in behavioural phenotype. Further longitudinal sampling and controlled experimental approaches will be needed to test these hypotheses and clarify the relative contribution of microbiome stability, environmental pressure, and individual plasticity.

The development of bacterial bioindicator indices provides a practical ecological tool for assigning individuals to behavioural clusters; here we have reached accuracies ranging from 75% to 89%. These indices may reduce the need for long-term behavioural monitoring by allowing behavioural tendencies to be inferred from a single faecal sample. This approach is particularly relevant for conservation programmes, where non-invasive and rapid tools could be needed to support individual assessment, welfare monitoring, and reintroduction planning. In applied terms, these bioindicator indices may help identify individuals that are more likely to belong to specific behavioural clusters. For example, assignment to the AB cluster could indicate a higher likelihood of poor welfare, health imbalance, or exposure to stressful environmental conditions. By contrast, assignment to the SB or LS clusters may suggest behavioural profiles with lower levels of anomalous behaviour, and therefore potentially less harmful phenotypes, given the known relevance of stereotypic behaviours as welfare indicators in captive animals (Lewis et al., 2022; Clive J.C. Phillips et al.,2017; Vanitha Varadharajan et al., 2016). Such information could support decisions about individual management, environmental enrichment, and release suitability. In addition, behavioural classification based on faecal biomarkers may contribute to predicting how lynxes interact with their territory after release, including their activity level and potential movement patterns. Although these indices should not replace direct behavioural observations, they offer a complementary, non-invasive tool for improving conservation management in both captive and reintroduced populations. Beyond their diagnostic value, these bioindicator indices offer considerable potential for individual-based management within conservation programmes. By enabling rapid behavioural classification, they may support the early identification of individuals requiring targeted interventions, such as environmental enrichment, health monitoring, or behavioural conditioning prior to release. Furthermore, their application could facilitate adaptive management strategies by allowing caretakers to tailor conditions and decisions according to the inferred behavioural profile of each individual

The lived-in environment strongly shaped gut microbiome structure and is closely associated with behaviour (Diaz & Reese, 2021). The Bayesian mediation model supported this relationship by showing that breeding station influenced both gut microbiome composition and behavioural phenotype. These results indicate that environmental context can affect behaviour directly while also modifying the gut microbiome in ways that may contribute to behavioural variation (Cryan et al., 2019; Matsuzaki et al., 2023).. The relative importance of these pathways differed among behavioural phenotypes. For anomalous behaviour, the strongest effect appeared to come directly from the environment, suggesting that external conditions may be especially relevant for the expression of stress-related or stereotypic behaviours. By contrast, sedentary behaviour and locomotion-and-surveillance showed a more balanced pattern, with both direct environmental effects and indirect effects mediated through the gut microbiome contributing to the observed behavioural profiles. Overall, the mediation model suggests that the environment and gut microbiome are interconnected components influencing behaviour rather than independent factors as it is said in Cryan et al (2019). However, the microbiome does not appear to be merely a passive reflection of the surrounding environment; instead, it may actively modulate host responses, potentially amplifying or buffering environmental pressures on behavioural expression (Diaz & Reese, 2021).

Building on the findings of Mortelliti et al. (2020), we propose that the gut microbiome can be interpreted as an ecosystem in which microbial heterogeneity may expand the range of behavioural phenotypes expressed by the host. In this framework, a diverse microbial community may operate similarly to a heterogeneous landscape, providing a broader set of biological conditions that support behavioural variability. Conversely, a more homogeneous microbiome may constrain this variability, favouring a narrower range of behavioural expressions. This perspective shifts away from the simplistic classification of microbiota as “beneficial” or “harmful”, and instead emphasizes its role in modulating phenotypic diversity. Our results provide empirical support for this idea. The LS cluster consistently showed the highest values across all diversity indices, a pattern that parallels the differences observed between breeding stations, where Granadilla exhibited higher microbial diversity than Acebuche. In particular, the strongest difference was found in the Shannon index between SB and LS, which is consistent with previous studies linking higher microbial diversity to more exploratory behaviour in birds (Florkowski et al., 2023). Together, these findings suggest that higher microbial diversity may be associated with more active or exploratory behavioural profiles. Furthermore, global diversity analyses revealed clear differences between LS and both AB and SB clusters, whereas no significant differences were detected between AB and SB. This pattern is consistent with the dendrogram structure, which showed that AB and SB are more closely related to each other than to LS. Taken together, both behavioural and microbiological analyses converge in identifying the LS cluster as the most distinct phenotype. However, these results should be interpreted with caution. Global diversity metrics alone may not fully capture the functional interactions between the microbiome and the host, as highlighted in previous studies (Reese et al., 2018; Shanahan, 2023; Williams et al., 2024). Therefore, although microbial diversity appears to be associated with behavioural variability, a more comprehensive understanding requires considering taxonomic composition and functional potential alongside diversity metrics.

The shifts on bacterial taxa and predicted functional pathways may provide a clearer understanding of the mechanisms linking gut microbiome composition with behavioural variation (Johnson et al., 2020; Pfau et al., 2023; Williams et al., 2024). In our study, gut microbiota composition differed appreciably across the three behavioural clusters, with distinct differential abundance patterns for key genera.The AB cluster shows a determinate microbial profile with relevant biological and behavioural implications. The higher abundance of Negativibacillus and Faecalibacterium observed in this cluster has previously been associated with neurodevelopmental and metabolic alterations, including depression, hyperinsulinemia, and autism (Zhang et al., 2025; Shi, et al.,; Iglesias-Vázquez et al., 2020). Conversely, the reduced abundance of Lachnospiraceae GCA-900066575 in this cluster is noteworthy, as decreases in this genus have been linked to impaired individual development (Bo et al.,2022). The microbial composition of the AB cluster could promote unusual neurological or metabolic states, caused by key microbial genera that have been consistently associated with abnormal conditions in the literature. Importantly, these microbial patterns align with behavioural characterisation of AB lynxes, which have a number of negative behavioural indicators.

The microbial composition of the SB cluster seems to be closely associated with decreased activity levels. Consistent with previous research, the higher prevalence of the [Ruminococcus] gnavus group, [Ruminococcus] gauvreauii group, and Allobaculum in this cluster is in line with findings associating these taxa with longer periods of sleeping and a more sedentary lifestyle (Chuang, H. H., 2024; Yuan et al., 2022; Watanabe et al.,2023; Chen et al.,2026), while a few studies have discovered weaker or less reliable links between these genera and depressive traits (Wei et al., 2024). Genera *UCG-005*, *Parabacteroides*, and *Prevotella_7* were less abundant in the SB cluster than in the LS and AB clusters. The reduced presence of these taxa is notable given their reported associations with lower exercise levels (Shi Y., 2022; Li Y., 2023) and their ambiguous or context-dependent associations with neurodevelopmental conditions such as autism (Iglesias-Vázquez L., 2020; Strati F., 2017).

Also, the LS cluster display a characteristic microbial profile. Genera more abundant in this cluster —including Peptoniphilus, NK4A214 group, Coprococcus, Staphylococcus, Enterococcus, Sphingobacterium, and Rikenellaceae RC9 gut group— have been linked to decreased anxiety and stress in human studies, the absence of traits related to autism, increased social behaviour (Liang et al., 2022; He et al., 2024; Levkova et al., 2023; Wu et al., 2021), and, in some cases, precocious puberty (Rodríguez Mazariegos et al., 2025). Nevertheless, the literature also reports associations between elevated levels of Coprococcus and Sphingobacterium in conditions such as obesity (Pisanu et al., 2020) and NK4A214 group in attention deficit hyperactivity disorder (Tang K., 2022). The association of Rikenellaceae RC9 gut group with stress reported in previous studies (Zhang et al., 2023) was not strongly supported by our data, suggesting that its functional role may vary across hosts or ecological contexts (Yurkovetskiy et al., 2015). Despite the amount of literature on the neural-gut microbiome theme, there is no data for Iberian Lynx, so it is important to acknowledge that most of these effects are probably unique to individual species. Extrapolating findings from one species to another requires exercising caution due to the added complexity (Yurkovetskiy et al., 2015). The absence of established associations limits interpretation but also highlights potential avenues for future investigation. In this study, the overall pattern observed in the LS cluster was more closely with favourable behavioural indicators, including low stress and an active lifestyle.

Functional profiling revealed cluster-specific metabolic pathways that may contribute to behavioural phenotypes. The AB cluster was enriched in pathways potentially associated with negative physiological effects, including the superpathway of lipopolysaccharide biosynthesis (Wen et al., 2022) and L-tryptophan degradation XII, which produces kynurenine, a precursor linked to neurotoxic effects (Agus A, 2018). In contrast, pathways associated with the LS cluster were mainly related to metabolic processes with potential roles in host physiological regulation and neural function. These included the L-methionine salvage cycle III and S-methyl-5-thio-α-D-ribose 1-phosphate degradation, both linked to S-adenosyl-L-methionine metabolism (Jeong et al., 2022), as well as pathways involving 2-oxoglutarate, associated with neural function (Tomé D, 2018; Qi W, 2025). Additionally, pathways related to mevalonate biosynthesis and isoprenoid production, which are important for cellular homeostasis (Gough, E. K, 2025), were more abundant in this cluster. Finally, peptidoglycan biosynthesis pathways, known to interact with host immune and neural systems and associated with social behaviour, were also enriched (Cryan et al., 2019). Overall, these results indicate clear functional differences between behavioural clusters, with AB enriched in potentially detrimental pathways and LS associated with pathways linked to physiological balance and host–microbiome interactions.

## Conclusion

Overall, this work provides a multi-layered framework linking behaviour, gut microbiota, and environmental factors in a critically endangered carnivore. By combining behavioural clustering, microbial community analysis, functional prediction, biomarker development, and mediation modelling, the study advances understanding of how captive environments may influence individual phenotypes.

