## Supporting information for "Unveiling correlational nexus among environment, gut microbiota, and personality traits in the Iberian Lynx (*Lynx pardinus*)"

[1]Ecología del Microbioma Global. Centre d'Estudis Avançats de Blanes (CEAB-CSIC)

[2]Centro de cría del lince ibérico El Acebuche (Huelva). Organismo Autónomo de Parques Nacionales (OAPN-MITECO)

[3]Centro de cría del lince ibérico Zarza de Granadilla (Cáceres). Organismo Autónomo de Parques Nacionales (OAPN-MITECO)10

Funding: project: INTERACTOMA, Agencia Estatal de Investigación (AEI) y fondos FEDER

**Figure captions:**

**Figure S1. Hierarchical classification of Iberian lynx behaviours**. Behavioural categories are organized hierarchically, with outer labels representing individual behaviours and inner labels corresponding to broader behavioural categories. Behaviours related to the breeding season, together with positive, negative, neutral and social interactions, were excluded from subsequent behavioural analyses.

**Figure S2. Nearest Sequenced Taxon Index (NSTI) evaluation**. Left panel shows the Mantel correlation between filtered and original microbial community compositions across different NSTI thresholds. The central panel shows the number of ZOTUs retained after applying each threshold, whereas the right panel displays the cumulative mean relative abundance of retained ZOTUs. The dashed vertical line indicates the selected threshold (NSTI = 0.20).

**Figure S3**. **Metadata overview of the studied individuals**. (A) Graphical classification of individuals according to metadata variables. (B) Heatmap showing the frequency of medication administration for each individual.

**Figure S4**. **Seasonal climatic differences between breeding stations**. Mean seasonal values are shown for climatic variables recorded at each breeding station: high (cvh) and low (cvl) vegetation cover (dimensionless, 0–1); 2-m, dew point temperature (d2m, Kelvin); evaporation (e, m water equivalent); 2-m, air temperature (t2m, Kelvin); total cloud cover (tcc, dimensionless, 0–1); total precipitation (tp, m); and 10-m zonal (u10) and meridional (v10) wind components ($m\ s^{-1}$). Asterisks indicate significant differences between breeding stations ($P < 0.05$).

**Figure S5**. **Alpha- and beta-diversity metrics across behavioural clusters and breeding stations.** Boxplots showing the distance to centroid (betadisper), observed richness and Shannon diversity index according to behavioural cluster (A) and breeding station (B).

**Figure S6**. **Differentially abundant bacterial taxa associated with breeding station**. (A) Relative abundance of ZOTUs exhibiting significant differences between breeding stations according to ANCOM-BC2. (B) Heatmap summarizing ANCOM-BC2 pattern analysis. Cell colour indicates the estimated log-fold change, with purple denoting decreased abundance and orange denoting increased abundance. Taxonomic assignments (phylum, class, genus and ZOTU) are provided for each row.

**Figure S7**. **ANCOM-BC2 pattern analysis of bacterial genera associated with behavioural clusters**. Columns represent pairwise comparisons among behavioural clusters, whereas rows correspond to bacterial genera identified as significantly differentially abundant. Cell colour represents the estimated log-fold change, with purple indicating lower abundance and orange indicating higher abundance relative to the reference group (AB cluster). Log-fold change estimates are displayed within each cell. Genera shown in black were significant before, but not after, sensitivity analysis, whereas genera highlighted in green remained significant after sensitivity analysis. Taxonomic information (phylum, class and genus) is provided for each row.

Supplementary table 1: Functional association between metabolic pathways and behavioural clusters.

| Group | Pathway | Sum up | Name | SB - AB | LS - AB | LS - AB | Bibiografical relevanze |
|---|---|---|---|---|---|---|---|
| Degradation of amino acids and phosphate compounds | LEU-DEG2-PWY | SB, LS> AB | L-leucine degradation I | 0.54 | 0.73 | 0 | X |
| | PWY-6505 | SB, AB> LS | L-tryptophan degradation XII (Geobacillus) | 0 | -0.53 | -0.71 | ✓ |
| | PWY-7527 | LS > SB, AB | L-methionine salvage cycle III | 0 | 0.86 | 0.66 | ✓ |
| | PWY-4361 | LS > SB, AB | S-methyl-5-thio-α-D-ribose 1-phosphate degradation | 0 | 0.89 | 0.66 | ✓ |
| Degradation of aromatic compounds | P281-PWY | SB, LS> AB | 3-phenylpropanoate degradation | 0.7 | 1.08 | 0 | X |
| | PWY-6210 | SB, AB> LS | 2-aminophenol degradation | 0 | -0.91 | -0.83 | X |
| Carbohydrates metabolism | PWY-5941 | LS > SB, AB | glycogen degradation II | 0 | 0.54 | 0.52 | X |
| | SUCSYN-PWY | LS > SB, AB | sucrose biosynthesis I (from photosynthesis) | 0 | 0.56 | 0.66 | X |
| | PWY-7347 | LS > SB, AB | sucrose biosynthesis III | 0 | 0.58 | 0.67 | X |
| Lipids/lipopolysaccharide biosynthesis | PWY-922 | LS > SB, AB | mevalonate pathway I | 0 | 0.82 | 0.89 | ✓ |
| | PWY-5910 | LS > SB, AB | superpathway of geranylgeranyldiphosphate biosynthesis I (via mevalonate) | 0 | 0.81 | 0.88 | ✓ |
| | LPSSYN-PWY | AB > SB, LS | superpathway of lipopolysaccharide biosynthesis | -0.98 | -1.23 | 0 | ✓ |
| Biosynthesis of wall compounds | PWY-5265 | LS > SB, AB | peptidoglycan biosynthesis II (staphylococci) | 0 | 0.56 | 0.58 | ✓ |
| | PWY-6470 | LS > SB, AB | peptidoglycan biosynthesis V (β-lactam resistance) | 0 | 0.65 | 0.63 | ✓ |

SUPPLEMENTARY FIGURE 1:

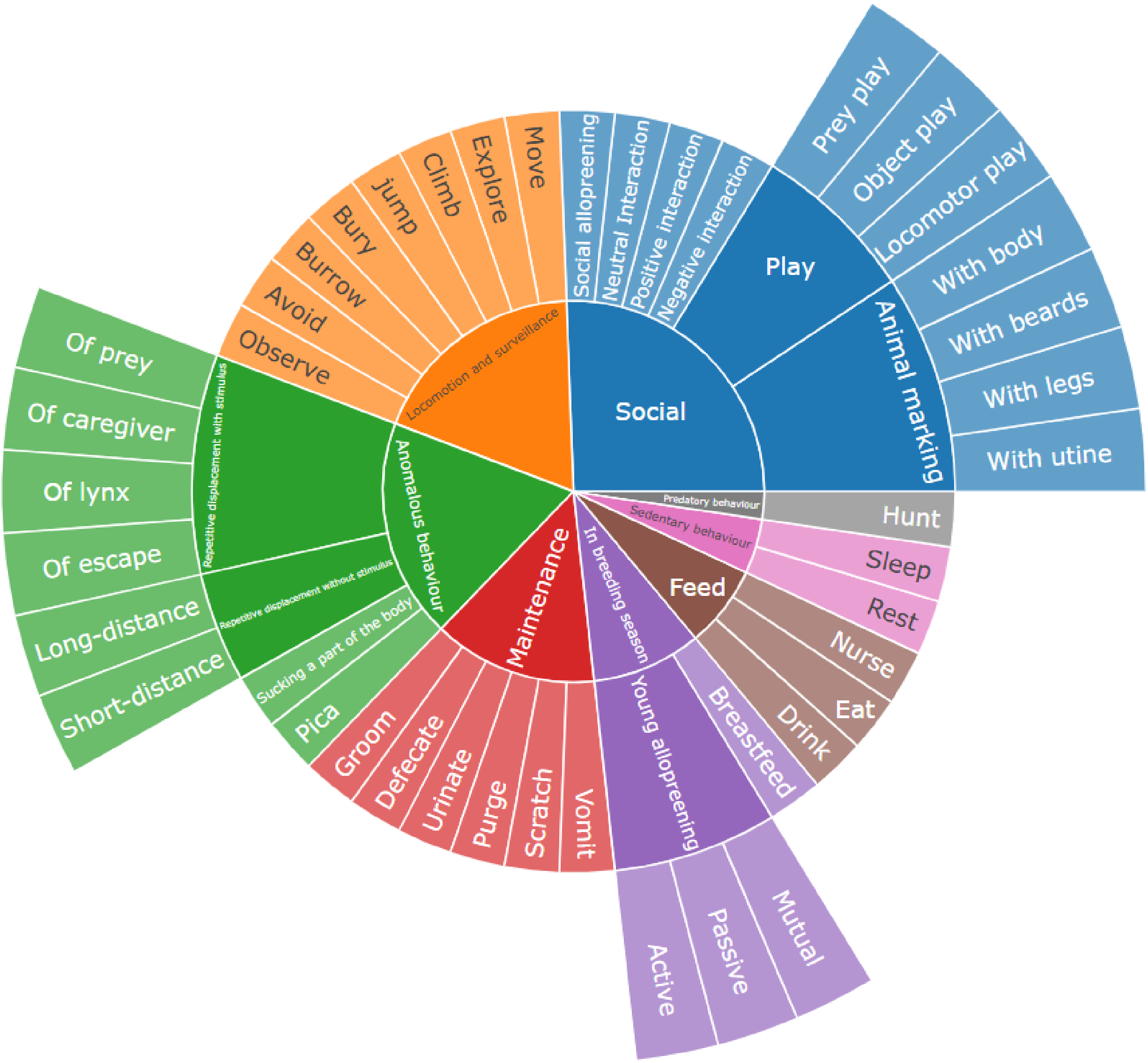

SUPPLEMENTARY FIGURE 2:

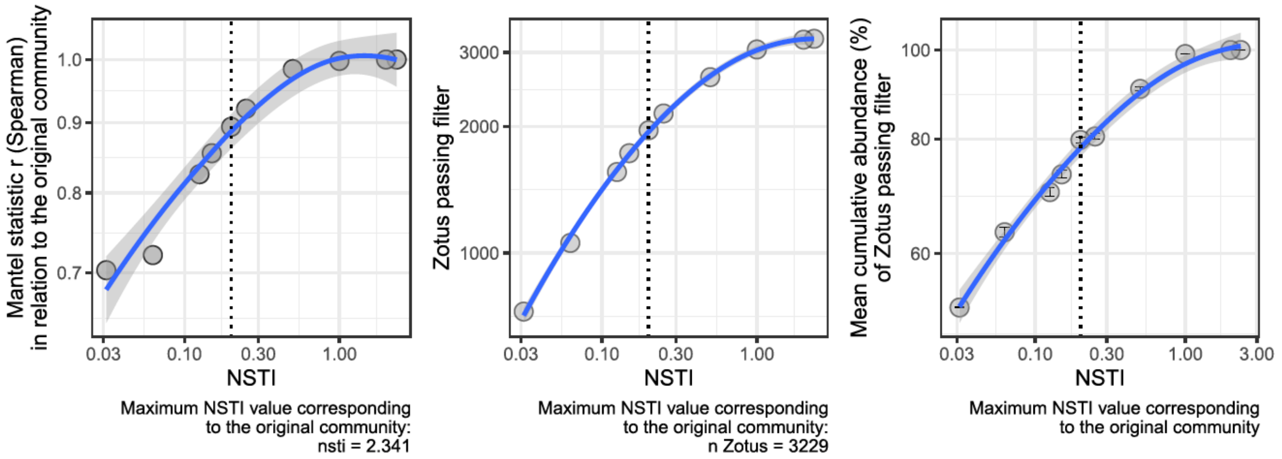

SUPPLEMENTARY FIGURE 3:

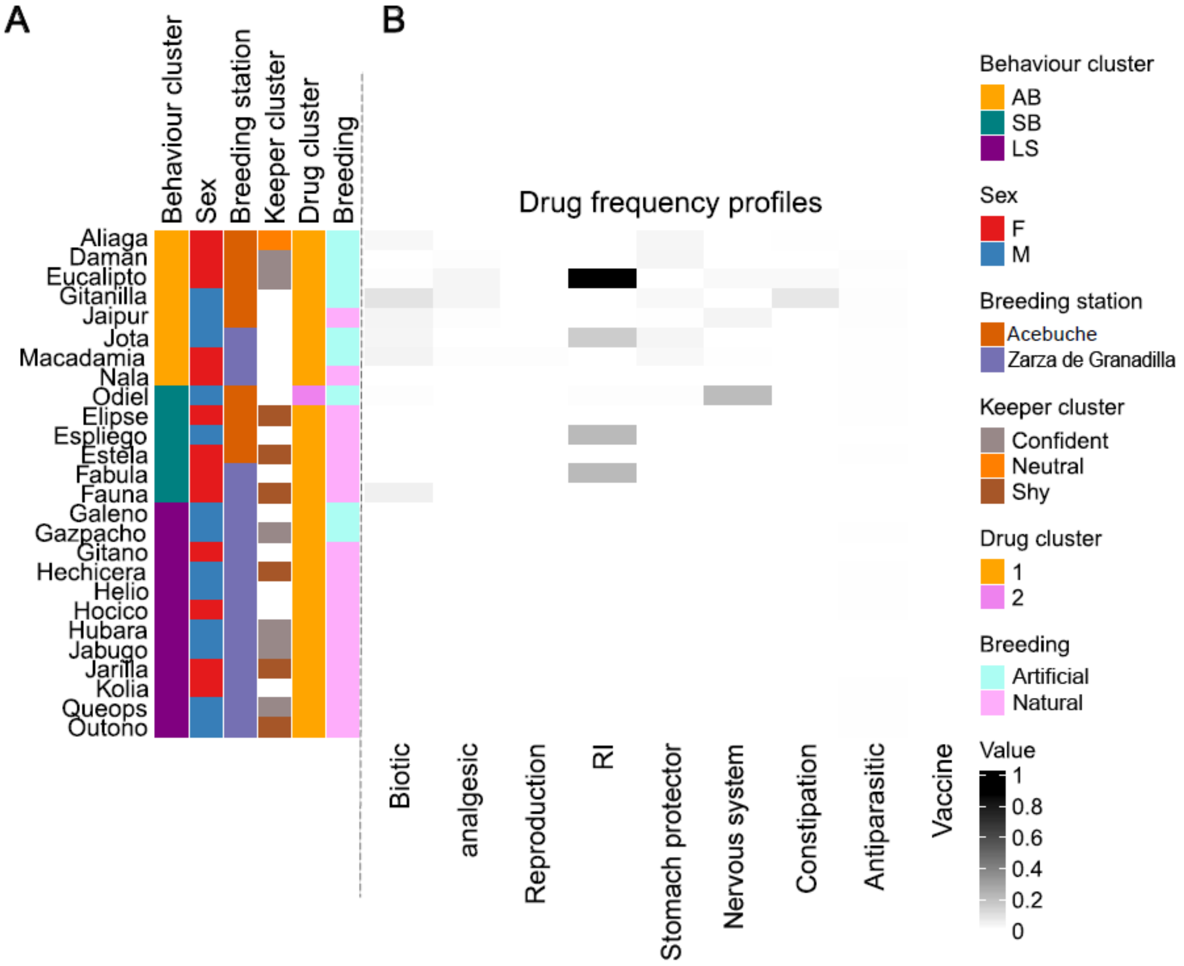

SUPPLEMENTARY FIGURE 4:

Seasonal evolution of climatic variables with significant differences

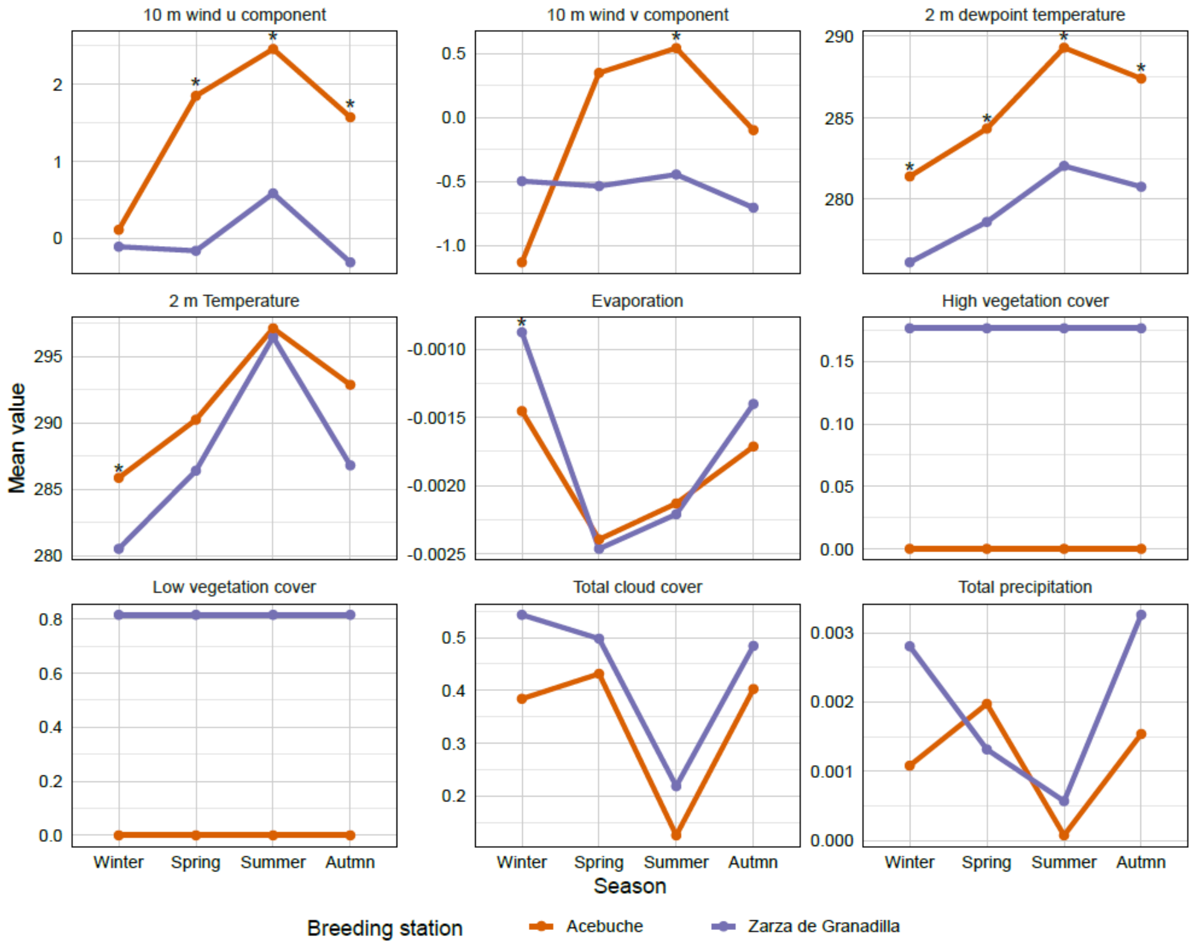

SUPPLEMENTARY FIGURE 5:

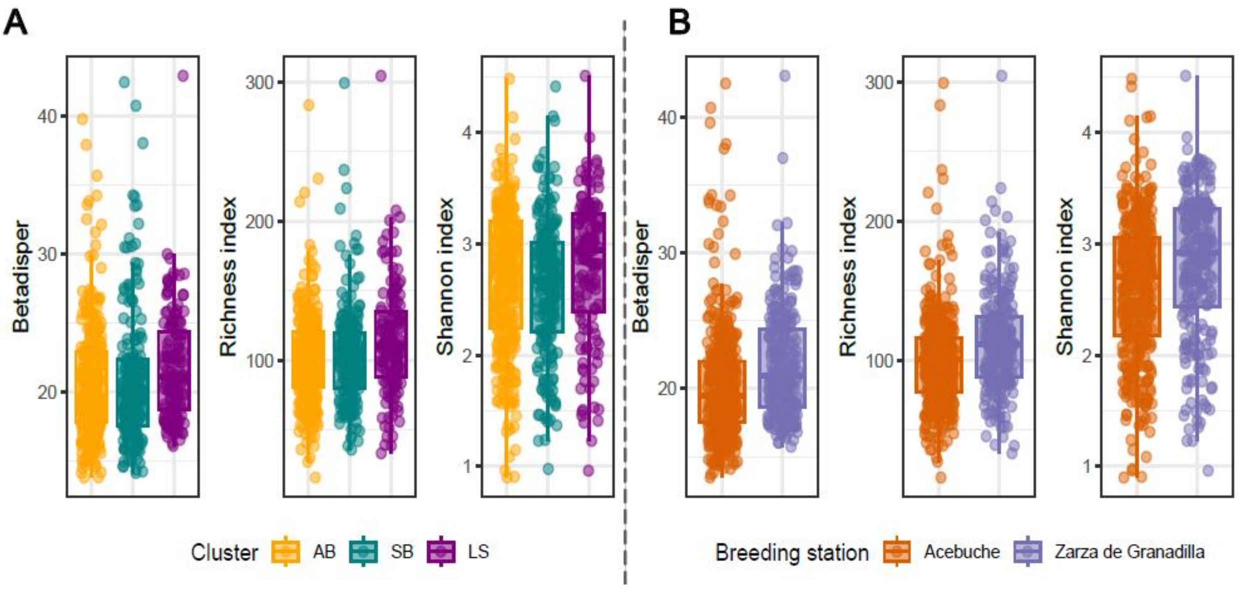

SUPPLEMENTARY FIGURE 6:

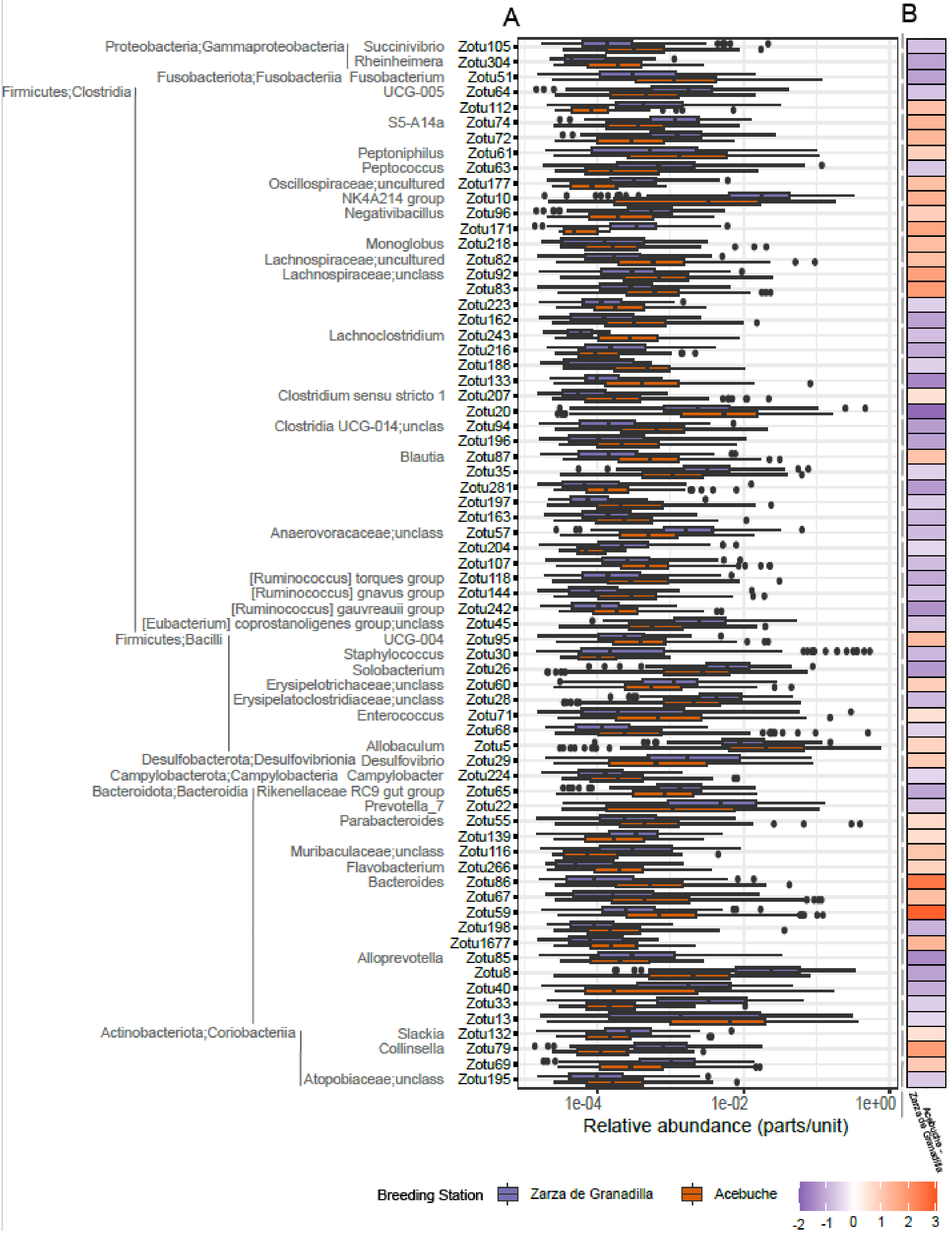

SUPPLEMENTARY FIGURE 7:

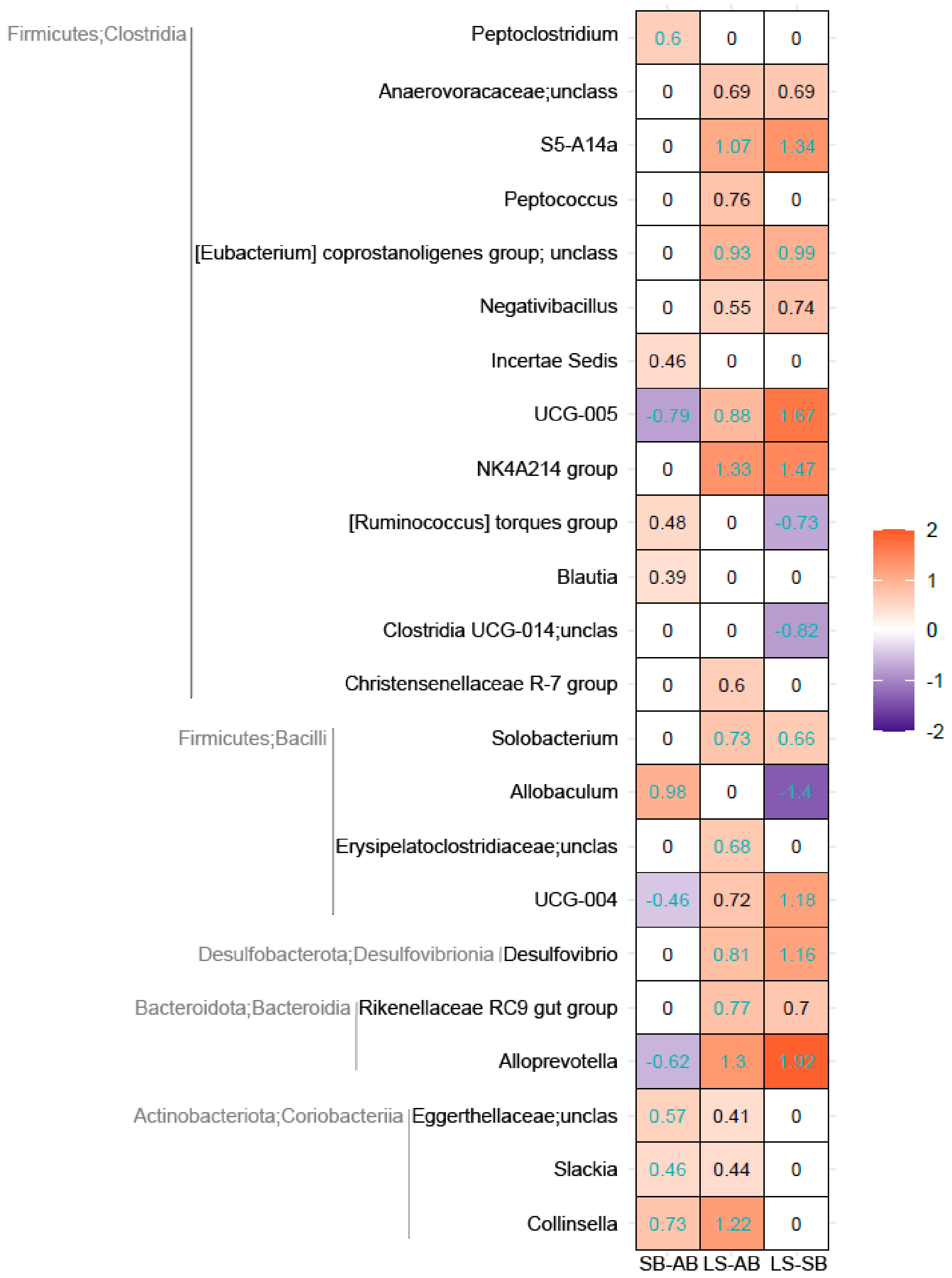
